# Extracurricular roles and divergent regulation of FAMA in *Brachypodium* and *Arabidopsis* stomatal development

**DOI:** 10.1101/2022.06.09.495393

**Authors:** Katelyn H. McKown, M. Ximena Anleu Gil, Andrea Mair, Shouling Xu, Dominique C. Bergmann

## Abstract

Stomata, cellular valves found on the surface of aerial plant tissues, present a paradigm for studying cell fate and patterning in plants. A highly conserved core set of related basic helix-loop-helix (bHLH) transcription factors (TFs) regulate aspects of stomatal cell identity in diverse species. We characterized BdFAMA in the temperate grass, *Brachypodium distachyon*, and found this late-acting TF was necessary and sufficient for specifying stomatal GC fate, and unexpectedly could also induce the recruitment of subsidiary cells in the absence of its paralogue, BdMUTE. The overlap in function is paralleled by an overlap in expression pattern and by unique regulatory relationships between BdMUTE and BdFAMA. To better appreciate the relationships among the Brachypodium stomatal bHLHs, we characterized the diversity of bHLH complexes through in vivo proteomics in developing leaves and found evidence for multiple shared interaction partners. The ability of BdFAMA to compensate for BdMUTE then prompted us to reexamine the roles of these paralogues in Arabidopsis. By testing genetic sufficiency within and across species, we found that, while BdFAMA and AtFAMA can rescue stomatal production in Arabidopsis *fama* and *mute* mutants, only AtFAMA can participate in the specification of Brassica-family-specific myrosin idioblasts. Taken together, our findings further our understanding of how the molecular framework of stomatal development has been reprogrammed across the monocot/dicot divide, as well as within the grasses, to produce distinct stomatal forms and patterns, and compel us to refine the current models of stomatal bHLH function and regulatory feedbacks amongst paralogues.

## INTRODUCTION

Cell fate establishment is essential for creating cellular diversity, for tissue development, and for organ formation—effectively all aspects of organismal development. Stomatal development represents a paradigm for studying cell fate acquisition following asymmetric cell divisions. Stomata, present in nearly all land plants, are cellular valves found on the surfaces of aerial plant tissues which regulate carbon dioxide and water vapor exchange. Stomata are typically made up of two kidney-shaped guard cells (GCs) flanking a pore, but the shape and patterning of these structures on a leaf can be clade-specific. The kidney-shaped GCs of dicots like *Arabidopsis thaliana* are distributed in pattern that follows a one-cell spacing rule, where little order exists beyond the prohibition of stomata in direct contact (Yang *et al*., 1995; Geisler *et al*., 2000). This pattern emerges from asymmetric and oriented divisions of dispersed stem cell-like precursors, called meristemoids, in the young leaf epidermis. Stomata in grasses have a distinctly different morphology, featuring dumbbell-shaped GCs flanked by a pair of subsidiary cells (SCs) derived from adjacent, lineally unrelated, cell files. Their ontogeny is also highly ordered, and the grass stomatal lineages develop sequentially from the base of a developing leaf in epidermal cell files adjacent to those which overlie veins (Fig. 1A).

**Figure 1.**
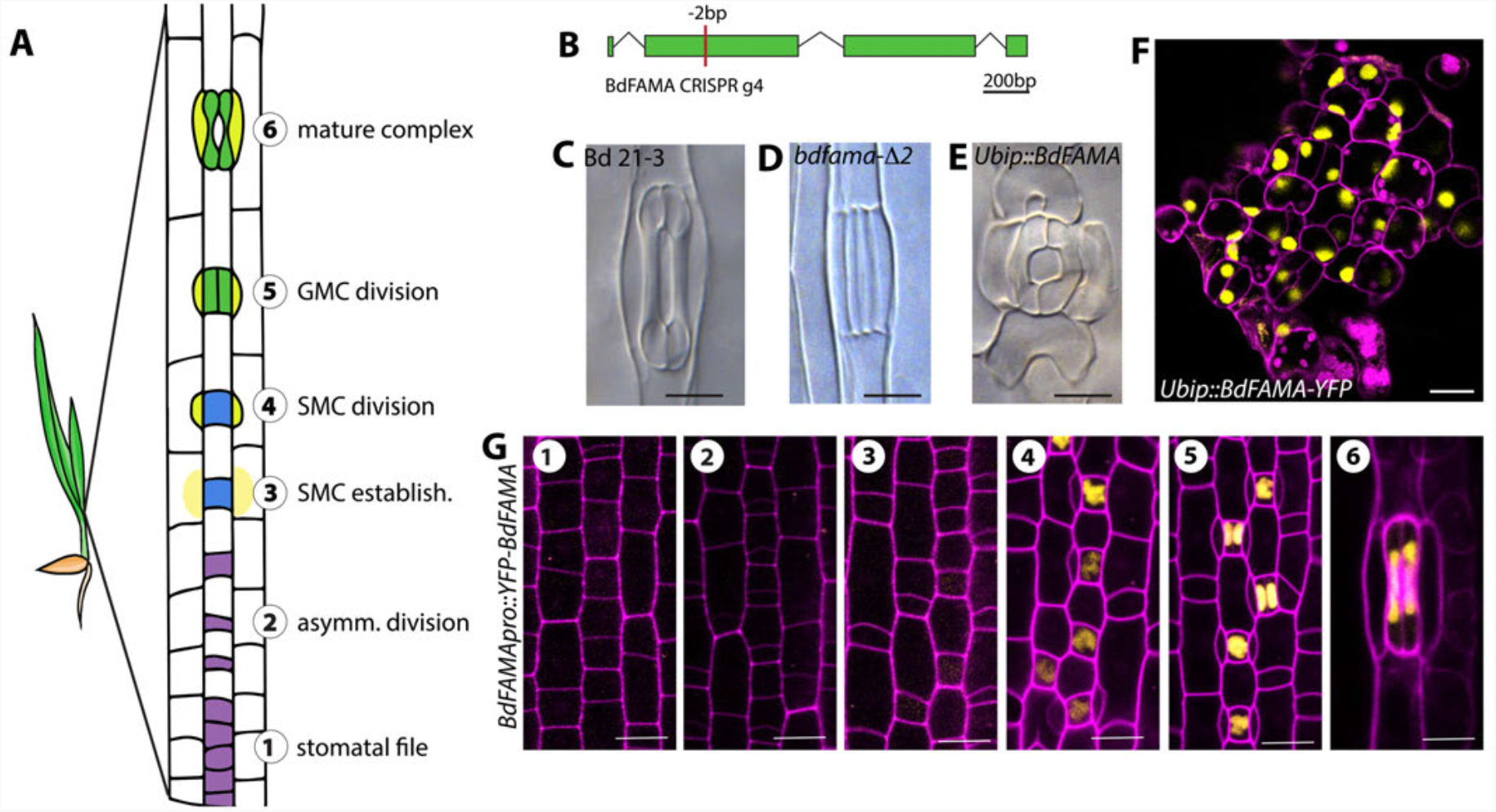
BdFAMA is required for stomatal guard cell differentiation. (**A**) Diagram of stomatal development in the developmental zone of a *Brachypodium distachyon* leaf, with stages 1-6 as follows. A stomatal file is established (purple) and the smaller daughter cell of an asymmetric division gives rise to a guard mother cell (GMC) (blue). GMCs induce subsidiary mother cell (SMC) formation from laterally adjacent cells, which divide asymmetrically to produce subsidiary cells from the smaller daughter cell (SCs) (yellow). GMCs divide symmetrically to form GC precursors (green), which differentiate to form a complete stomatal complex, with pairs of GCs and SCs flanking a pore. (**B**) Gene diagram of *BdFAMA* with location of CRISPR/Cas9-generated mutation site. (**C-E**) DIC microscopy images of stomata from cleared leaf tissue in WT (Bd21-3) **(C)** and *bdfama-*D*2* (**D**), and *Ubipro::BdFAMA* (**E**). **(F)** Confocal image of ectopic, unpaired guard cells induced by *Ubipro::BdFAMA-YFP* (yellow). **(G)** Confocal images of *BdFAMApro::YFP-BdFAMA* reporter expression during stomatal development. No *YFP-BdFAMA* expression is detected during stomatal file establishment and asymmetric division (stages 1 and 2). *YFP-BdFAMA* (yellow) is detected at low levels in GMCs during SMC establishment (stage 3), and increases in expression during SC recruitment (stage 4). *YFP-BdFAMA* peaks in newly formed GCs (stage 5), and persists in mature stomatal complexes (stage 6). DIC images taken of cleared abaxial epidermal tissue; images of *bdfama-*D*2* were from the 2^nd^ leaf of 11dpg T4 plants, *Ubipro::BdFAMA* and *Ubipro::BdFAMA*-YFP are from T0 regenerants. Confocal images from 2^nd^ leaf 6dpg plants, cell walls (magenta) stained with Propidium Iodide (PI). Scale bars: 10 μm. **See also Figure S1**

*Arabidopsis FAMA* (*AtFAMA*) was the first transcriptional regulator published as being essential for stomatal fate (Bergmann *et al*., 2004; Ohashi-Ito *et al*., 2006), and multiple studies since have linked this factor to both the acquisition and the continued maintenance of the differentiated state, the latter through direct interaction with cell cycle regulator, RETINOBLASTOMA RELATED (RBR) (Hachez *et al*., 2011; Lee *et al*., 2014; Matos *et al*., 2014). *AtFAMA* has additional roles in cell cycle control; in *atfama* mutants, epidermal tumors form by the repeated symmetric cell divisions of incompletely differentiated guard cells (Ohashi-Ito *et al*., 2006). *AtFAMA* and its two closest paralogues, *SPEECHLESS* (*AtSPCH*) and *AtMUTE*, are basic helix-loop-helix (bHLH) transcription factors in the group Ia subfamily (Pires & Dolan, 2010), and together they initiate the asymmetric divisions of the stomatal lineage and ensure commitment to guard cell fate, respectively (Ohashi-Ito *et al*., 2006; Pillitteri *et al*., 2007; MacAlister *et al*., 2007). Expression of *AtFAMA* can produce single cells with hallmarks of stomatal GC identity in the absence of *AtSPCH* and *AtMUTE*, but hyperactivation of these earlier-acting TFs cannot overcome the requirement for *AtFAMA* to produce GCs (Ohashi-Ito *et al*., 2006). AtFAMA is also required as part of a negative regulatory loop with AtMUTE to ensure that stomata consist of exactly one pair of sister guard cells (Han *et al*., 2018). To bind DNA and effect transcriptional change, AtSPCH, AtMUTE, and AtFAMA require heterodimerization with functionally redundant group IIIb bHLH partners, INDUCER OF CBF EXPRESSION1/SCREAM (AtICE1/AtSCRM) and AtSCRM2 (Kanaoka et al., 2008).

The genetic origin and duplication history of SPCH, MUTE, FAMA, ICE1/SCRM and SCRM2 has been tracked among extant land plants, and there is evidence for deep conservation and connection to stomatal identity (Chater *et al*., 2016). Among flowering plants, orthologues of *AtFAMA* are readily identified (Macalister & Bergmann, 2011; Chater *et al*., 2016; Raissig *et al*., 2016; Wu *et al*., 2019; Ortega *et al*., 2019), and preliminary functional analysis of mutants in rice indicate that *FAMA* orthologs are necessary for the production of functional GCs, though not for cell cycle inhibition, as *osfama* mutants arrest with two incompletely differentiated GCs and SCs (Liu *et al*., 2009; Wu *et al*., 2019).

Other bHLHs have undergone clade-specific gene duplications. For example, compared to *Arabidopsis*, the temperate grass model, *Brachypodium distachyon*, (Pooidae) has an additional *SPCH*, and these two *BdSPCH* paralogues have partially redundant roles in lineage initiation. Both *Arabidopsis* and *Brachypodium* have two SCRM-like genes, but they are derived from independent duplications, and they exhibit different degrees of functional redundancy in stomatal lineage activities (Kanaoka *et al*., 2008; Raissig *et al*., 2016). Recent data has also shed light on functional divergences within the grasses. The single *MUTE* gene which in grasses acquired a new role in SC recruitment, is essential in maize and rice, but not in *Brachypodium* (Raissig *et al*., 2017; Wu *et al*., 2019; Wang *et al*., 2019). The differential requirement for *MUTE* to produce viable stomata suggests specialization can occur, even within this well-conserved gene family, and such specialization may extend to other elements of the stomatal lineage developmental regulatory network.

Of the key players in stomatal development, *SPCH1* and *SPCH2, MUTE, SCRM2*, and *ICE1* have all been characterized in *Brachypodium*, but the expression and function of the final group 1a bHLH, *FAMA*, has not. Evidence from phylogenetic and sequence homology support *FAMA* as being closest to an ‘ancestral’ form from which the group Ia paralogs were derived (Macalister & Bergmann, 2011), and unlike SPCH and MUTE, the FAMA function of GC differentiation is completely dependent on an intact DNA binding domain (Davies & Bergmann, 2014). Although the GC differentiation role of *FAMA* is conserved across land plants, FAMA has demonstrated additional functions in *Arabidopsis* such as cell division control and enforcement of terminal cell fate through chromatin remodeling, and it has been co-opted into the development of a Brassica-specific non-epidermal cell type. Whether the roles of FAMA extend beyond its GC differentiation capacity in other plant systems, however, is not known. In this study, we characterize the endogenous roles of *BdFAMA* and find that it necessary and sufficient for specifying stomatal GC fate. Our data show that *BdFAMA* expression commences much earlier in the stomatal lineage than expected given its terminal role in GC differentiation, and this extended expression may allow *BdFAMA* to compensate for the absence of *BdMUTE* in a way that is distinct from the other grasses. Because loss of *BdFAMA* resembles the loss of *BdSCRM2*, and BdFAMA and BdMUTE have overlapping functions, we characterized the diversity of bHLH complexes through *in vivo* proteomics in developing *Brachypodium* leaves and found evidence for multiple shared interaction partners. The overlaps in *BdFAMA* and *BdMUTE* function then prompted us to reexamine the roles of these paralogues in *Arabidopsis*. By testing genetic sufficiency within and across species, we found that *BdFAMA* and *AtFAMA* can rescue stomatal production in *Arabidopsis fama* and *mute* mutants, but that only *AtFAMA* can participate in the specification of Brassica-family specific myrosin idioblasts. Taken together, our findings prompt us to reconsider the current models of stomatal bHLH function and enabled us to refine our understanding of regulatory feedbacks amongst paralogues.

## RESULTS

### Functional analysis of *BdFAMA*

To characterize the role of the putative *Brachypodium FAMA* homolog (BdFAMA, Bradi2g22810) in stomatal development, we used CRISPR-Cas9 genome editing in the wildtype line, Bd21-3, to induce coding region mutations early in the second exon. As expected from the essential nature of *FAMA* in other species, T0 regenerants harboring homozygous frameshift mutations were seedling lethal. We were able to obtain a transgenic plant that was heteroallelic for a 6 basepair (bp) and a 2bp deletion, with the latter predicted to result in a frameshift and an early stop in the second exon, prior to the bHLH domain (Fig. 1B and Fig. S1A-C). Among offspring of this line, plants homozygous for *bdfama-*Δ*2* completely lacked functional stomata, and instead exhibit a four-celled complex of undifferentiated, paired GCs, with flanking subsidiary cells (SCs) (Fig. 1C-D). The same stomatal defective phenotype was observed in T0 regenerants bearing other homozygous frameshift mutations (Fig. S1D). In contrast to the *fama* ‘tumors’ found in *Arabidopsis*, there were no continued GMC divisions in *bdfama*, which supports a role for *BdFAMA* in GC differentiation, but not necessarily cell division control. The undifferentiated four-cell complex phenotype is strikingly similar to the phenotype resulting from a loss of BdFAMA’s predicted partner *BdSCRM2*, but differs from that seen in from loss of the other potential heterodimer partner *BdICE1* (Raissig *et al*., 2016).

Loss of *BdFAMA* resulted in a discrete phenotype of GC differentiation failure without disturbing general leaf shape and pattern, but broad overexpression of *BdFAMA* (*Ubipro:BdFAMA-YFP* and *Ubipro::BdFAMA*) resulted in severely deformed leaves where most leaf epidermal cells were converted into kidney-shaped cells that accumulated localized cell wall material resembling a pore (Fig. 1E-F, Fig. S1E). The vast majority of these cells were unpaired and oriented such that their pore-like region faced the leaf tip, a phenotype that resembles the overexpression of AtFAMA in *Arabidopsis* (Ohashi-Ito *et al*., 2006). Occasionally, paired GCs with flanking cells resembling SCs were seen near files adjacent to cells overlying veins (Fig. 1E), which may suggest that cells already fated to become GCs retain some normal patterning, whereas non-stomatal cells transdifferentiate when exposed to BdFAMA. Taken together, these results suggest *BdFAMA* is necessary and sufficient to promote GC fate.

To monitor BdFAMA expression in developing leaves, we created transcriptional (*BdFAMApro::3xYFP*) and translational (*BdFAMApro::YFP-BdFAMA*) reporters. We anticipated that *BdFAMA* would be restricted to the final stage of stomatal development, consistent with the loss of terminal GC differentiation in *bdfama. BdFAMApro::YFP-BdFAMA*, however, could already be detected weakly in young GMCs during the timeframe of SMC establishment. This reporter peaked before the GC division, and persisted into mature GCs (Fig. 1G). A similar pattern was seen with the transcriptional reporter (Fig. S1F). The BdFAMA expression window overlaps considerably with that of *BdMUTE* (Raissig *et al*., 2017) during SC recruitment, GC division, and early GC differentiation; although unlike *BdMUTE, BdFAMA* does not appear to be expressed in SMCs or SCs. *BdFAMA*’s temporally broad and continued expression contrasts with the situation in *Arabidopsis*, where AtFAMA peaks in young GCs, and both translational reporters (Adrian *et al*., 2015; Han *et al*., 2018) and single cell RNA sequencing (scRNAseq) data (Lopez-Anido *et al*., 2021) show *AtMUTE* and *AtFAMA* expression initiates sequentially, and that they share little overlap in protein expression.

### *BdMUTE* is involved in regulating *BdFAMA* during development

In *bdmute*, the predominant phenotype is failure to recruit SCs, such that stomatal complexes consist solely of a pore flanked by two GCs with kidney-rather than dumbbell-shaped morphologies (Raissig *et al*., 2017). In addition, 40% of the complexes also show defects in GC differentiation, and instead produce long, paired cells that resemble the undifferentiated GCs of the *bdfama* mutant. In the mature leaf zone of *bdmute*, these paired cells never take on GC fate, and instead appear to leave the stomatal lineage, elongating in a way similar to pavement cells. Based on these phenotypes, we hypothesized that *BdMUTE* is involved in the expression or regulation of *BdFAMA*, and that the aborted GCs were a result of insufficient BdFAMA activity. To test regulatory relationships between *BdMUTE* and *BdFAMA*, we created transgenic lines where the *BdFAMA* translational reporter was expressed in a *bdmute* background (Fig. 2 A-B). We detected YFP-BdFAMA in all mature, differentiated GCs in *bdmute; BdFAMApro::YFP-BdFAMA;* however, we did not observe YFP signal in the undifferentiated complexes, which suggests that the failure to differentiate was indeed due to lack of BdFAMA. It is unclear whether the absence of *YFP-BdFAMA* in aborted GCs is due to complete failure to activate *BdFAMA* expression in stomatal precursors or a failure to maintain sufficient levels of BdFAMA, but it does suggest that *BdMUTE* likely plays a role in stabilization of *BdFAMA* expression during development.

**Figure 2.**
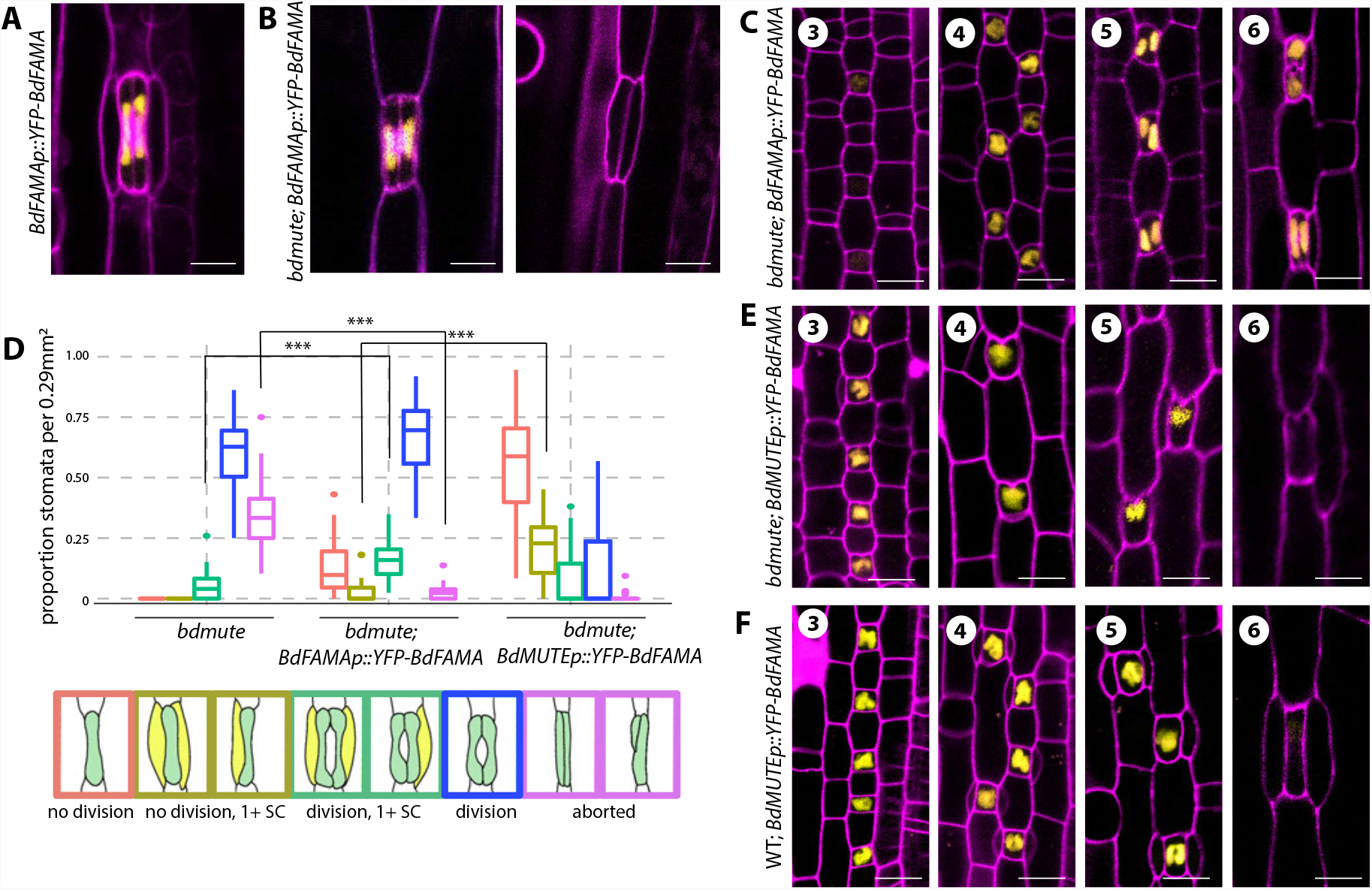
BdFAMA is able to recruit subsidiary cells in the absence of *BdMUTE*. (**A-C**) Confocal images of *BdFAMApro::YFP-BdFAMA* in stomatal complexes of wild type **(A)** and *bdmute* **(B)**. *YFP-BdFAMA* is present in differentiated complexes but is absent in aborted GCs (**B**). **(C)** Confocal images of stomatal development in *bdmute; BdFAMApro::YFP-BdFAMA* during SMC establishment (stage 3), and SC recruitment (stage 4), newly formed GCs (stage 5), and mature stomatal complexes (stage 6). (**D**) Quantification of stomatal phenotypes in *bdmute* and *bdmute* expressing either *BdFAMApro::YFP-BdFAMA*, or *BdMUTEpro::YFP-BdFAMA* from cleared abaxial tissue of the 2^rd^ leaf, 6 to 7 days post germination (dpg). Cartoon representations of phenotypic classes are shown at the bottom, outlined in the colors represented in the plot. For each sample, five different regions of the leaf were imaged and quantified. n=6 for *bdmute* and *bdmute; BdFAMApro::YFP-BdFAMA*, and n=8 for *bdmute; BdMUTEpro::YFP-BdFAMA*. ***P<0.001 (based on Wilcoxon-rank sum test followed by Dunn’s multiple comparisons test). In each boxplot, the colored horizontal line indicates the median, upper and lower edges of the box (hinges) represent the upper and lower quartiles; whiskers extend to the largest observation within 1.5 interquartile ranges of the box. (**E-F**) Confocal images of stomatal development of *bdmute; BdMUTEpro::YFP-BdFAMA* and of wild type; *BdMUTEpro::YFP-BdFAMA* T0 regenerants at indicated stages. Cell outlines (magenta) visualized by Propidum Iodide (PI) staining. Scale bars: 10 μm. **See also Figures S2 and S3**

### *BdFAMA* is able to recruit subsidiary cells in *bdmute*

In surveying phenotypes in the multiple independent transformed lines of *bdmute; BdFAMApro::YFP-BdFAMA*, we noticed that many also exhibited a significant reduction in occurrence of aborted GCs. Using T4 plants from a representative line (line 19), we quantified this phenotype, and found that aborted GCs were reduced from ∼34% in *bdmute* to ∼2% in *bdmute; BdFAMApro::YFP-BdFAMA* (Fig. 2D, aborted). A new phenotype emerged where some GMCs completely failed to divide symmetrically, but the single *YFP-BdFAMA*-positive GMCs acquired a dumbbell shape, as if their GC differentiation program was intact, however, these single GCs did not form a pore (Fig. 2D, no division).

A second unexpected phenotype was the successful recruitment of SCs to stomatal complexes. About 19% of complexes, including those whose GMCs did or did not divide symmetrically, were able to recruit one or both SCs (Fig. 2C, 1+ SC categories). The presence of SCs suggests *BdFAMA* can partially compensate for *BdMUTE*. This ability of *BdFAMA* to compensate for loss of *BdMUTE* is especially interesting given that the cell-cell mobility of BdMUTE was hypothesized to be enable it to recruit SCs from cell files neighboring the GMC (Raissig *et al*., 2017). Our *BdFAMA* translational and transcriptional reporters were detected exclusively in GMCs in both *bdmute* and wildtype *Bd21-3* plants, so there is no evidence that the BdFAMA protein can move from cell to cell (Fig. 1G, Fig. 2C and Fig. S1F). While many of the complexes with successful SC recruitment had typical SCs—appropriately proportioned to the length and size of its stomatal complex—many SCs were abnormally large and arose from diagonal SMC divisions, and it was not clear whether these large SCs would be able to work in concert with the GCs in complex to facilitate rapid pore opening and closing (Fig. S2A). Many of the GMC divisions were also abnormal, and produced two asymmetric GC precursors (rather than two symmetric cells) or oriented their division planes perpendicular to the normal placement. The misorientation of GMC division occurs in both *bdmute* and *bdmute; BdFAMApro::YFP-BdFAMA*, so there does seem to be some requirement of *BdMUTE* for proper placement of division plane that is not improved with additional BdFAMA (Fig. S2B).

### Earlier expression of *BdFAMA* causes GMC division failure

While partial complementation of *bdmute* was possible with BdFAMA expressed under its own promoter, SC recruitment, when successful, seemed to occur later in development than it does in wildtype. To test whether rescue of SC recruitment in *bdmute* improved with earlier expression of BdFAMA, we created *BdMUTEpro::YFP-BdFAMA* in the *bdmute* background. Most T0 regenerants died as young plantlets, and in tissue imaged from these individuals (n=13 severe lines, n=6 milder lines), we observed that GMCs failed to divide symmetrically to produce paired GCs and instead produced single GCs with a dimple resembling a pore (pseudo-pores), oriented toward the leaf tip (Fig. 2E). These pseudo-pores also began forming in GMCs during the SC recruitment stage, indicative of premature BdFAMA activity (Fig. 2E, stage 3). Approximately 21% of these single GC complexes were able to recruit one or more SC (Fig. 2D). In some lines, single GC complexes were stacked end-to-end, violating the single cell spacing rule, and a large pore with underlying airspace was formed that allowed a stunted, pale green plant to persist and produce a few T1 seeds, which enabled us to quantify the relative expression levels across the lines (Fig. S3A-B). We observed a dose-dependent effect on GC division and plant growth; individuals in which GCs failed to divide symmetrically and caused lethality or stunted plant growth expressed more *YFP-BdFAMA* than lines with infrequent GC division phenotypes that had normal growth (Fig. S3C-E).

We hypothesized that the GMC division failure is caused by the premature activity of *BdFAMA* only in the absence of *BdMUTE*, and that normally BdMUTE acts as temporal placeholder that, through competition for binding partners or target promoter binding sites, ensures that differentiation does not commence until SCs are recruited and the GMCs have divided. To test this hypothesis, we expressed *BdFAMA* in the *BdMUTE* domain (*BdMUTEpro::YFP-BdFAMA)* in a wildtype background. We observed a dose-dependent effect; many T0 regenerants produced stomatal complexes with single GCs like in *bdmute; BdMUTEpro::YFP-BdFAMA* (n=7 severe lines). More lines (n=10 mild lines), however, had an intermediate or milder phenotype, where only some of their GCs failed to divide symmetrically. Additionally, if single GCs generated ectopic leaf-tip facing pores, they appeared only after SC recruitment (Fig. 2F). These data support a role for *BdMUTE* in the proper timing of *BdFAMA* activity.

### Isolation of stomatal bHLH complexes from *in vivo* co-immunoprecipitation with tagged BdMUTE and BdFAMA shows these proteins have shared and distinct interaction partners

How do *BdMUTE* and *BdFAMA* maintain their functional specificity with such overlap in expression and function? Based on similar mutant phenotypes, one might speculate that there is a preference for BdFAMA/BdSCRM2 heterodimers and for BdSPCHs/BdICE1 heterodimers (Raissig *et al*., 2016), but whether BdICE1 or BdSCRM2 preferentially heterodimerize with *BdMUTE* is not known. To better understand the functional overlap between BdMUTE and BdFAMA, we investigated *in vivo* binding partners using co-immunoprecipitation (Co-IP) and subsequent analysis with liquid chromatography with tandem mass spectrometry (LC-MS/MS). We sampled tissue from the proximal 4mm of leaves, a region that encompasses the peak expression of stomatal development genes from lines stably transformed with YFP-tagged BdSPCH2, BdMUTE, BdFAMA and BdSCRM2 expressed under their native promoters, and BdICE1, expressed under a Ubiquitin promoter (Fig. 3A and Fig. S4-5). A BdMUTE promoter-driven nuclear-targeted triple YFP-tagged line and wildtype Bd21-3 were used as controls.

**Figure 3.**
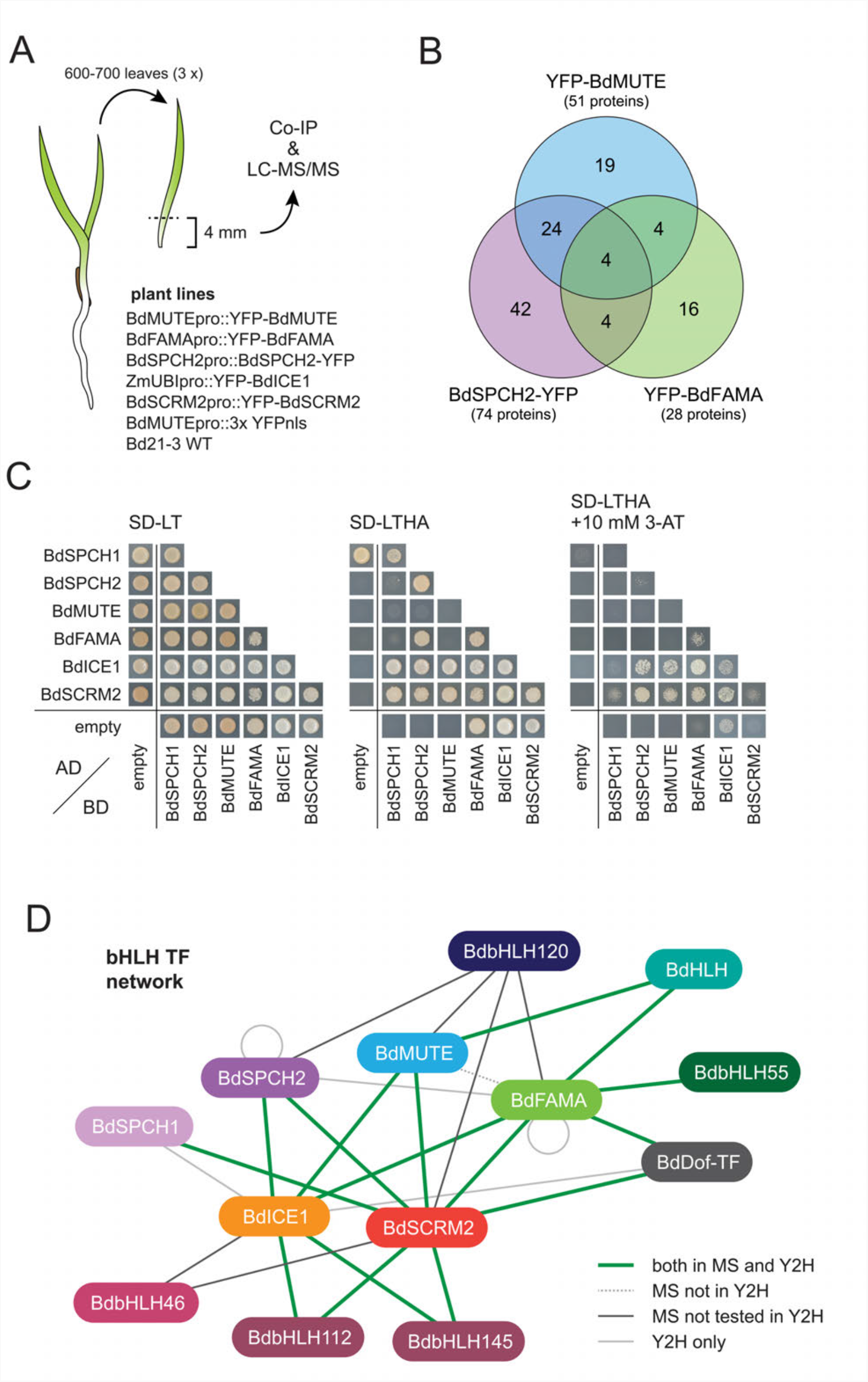
Identification of physical interaction partners of Brachypodium stomatal bHLHs. **(A)** Cartoon diagram of strategy and plant lines used to identify bHLH interactors *in vivo*. The developmental zones (bottom 4 mm) of 600-700 leaves per sample (3 replicates of each of 7 genotypes listed) were harvested and used for Co-IPs with GFP-Trap beads. Bound proteins were identified by LC-MS/MS. **(B)** Venn diagram showing the overlap of specific putative interaction partners found for BdMUTE, BdFAMA and BdSPCH2. **(C)** Y2H assay testing interactions between the six master regulators of stomatal development. Yeast transfected with the indicated AD-and BD-fusion proteins was spotted on SD-LT and SD-LTHA to test for successful transfection and protein interaction, respectively. 3-AT was added to increase stringency of the selection and overcome auto-activation **(D)** Stomatal bHLH TF network. Line styles indicate whether an interaction was identified in the Co-IP experiment (MS) and/or in the Y2H assays. **See also Figure S4-8**

With our model of preferred heterodimer pairing, BdFAMA/BdSCRM2 and BdSPCHs/BdICE1, we would expect to find these specific partner pairings from our pull-down interaction data. However, the LC-MS/MS data show that all assayed group Ia bHLHs, BdSPCH2, BdMUTE, and BdFAMA, are able to associate *in vivo* with both group IIIb bHLHs, BdICE1 and BdSCRM2. Despite possibly having shared heterodimer binding partners, we found that both BdMUTE and BdFAMA pulled down overlapping as well as unique interaction candidates; of the candidate interactors identified, only 8 were shared between BdFAMA and BdMUTE, which, based on our pipeline for defining enriched interactions, pulled down 28 and 51 total interactors, respectively (Fig. 3B, Fig. S6-7 and Table S3). Notably, the number of shared interactors between BdMUTE and BdFAMA is comparable to their overlap with BdSPCH2, suggesting shared interactors alone cannot explain the functional overlap of *BdFAMA* with *BdMUTE*. BdMUTE also pulled down BdFAMA, which suggested that the two might even exist in a shared complex. We also identified a number of transcription factors among interacting proteins, mainly bHLHs, that could act as alternative dimerization partners, or contribute to a higher order bHLH complex.

We confirmed these interactions *in vitro* with a yeast two hybrid (Y2H) assay and found that BdSPCH2, BdMUTE, and BdFAMA were able to interact with both BdICE1 and SCRM2 (Fig. 3C and Fig. S8). We could not confirm the interaction between BdMUTE and BdFAMA, and it is possible that these associate through a higher order complex rather than through direct interaction (Fig. 3C). We also confirmed several novel interactors. An HLH that might act as a transcriptional inhibitor interacted with BdMUTE and BdFAMA; bHLH55, for which an ortholog was found to interact with AtFAMA (Mair *et al*., 2019), interacted with BdFAMA; bHLH112 and bHLH145, which are orthologous to AtbHLH71, interacted with BdICE1 and BdSCRM2; and a Dof TF that showed similarity to STOMATAL CARPENTER (SCAP1), a GC morphology regulator in *Arabidopsis* (Negi *et al*., 2013) interacted with BdICE1 and BdSCRM2, and weakly with BdFAMA. The summary of core bHLH interaction data in Fig. 3D shows that although there is apparent flexibility of heterodimer pairings possible between bHLHs, unique interactions remain prevalent. Additionally, while there still might be preferred heterodimer binding partners that could influence functional specificity of the core stomatal bHLHs, it is not on a level that we were able to detect from these approaches. However, the possibility remains that BdFAMA could achieve the functional flexibility needed to substitute for *BdMUTE* by timing-or dosage-dependent alternate binding with BdICE1 or BdSCRM2.

### *BdFAMA* can drive the cell fate transition from GMC to GC in the absence of *BdMUTE*

In contrast to *Brachypodium*, maize and rice plants do not survive the loss of *MUTE* activity (Wu *et al*., 2019; Wang *et al*., 2019). GMCs in these cereal crops fail to recruit SCs and fail to divide longitudinally to form GCs, arresting instead as small cells of unclear identity after several misoriented divisions (Wu *et al*., 2019; Wang *et al*., 2019). Our observations that an extra copy of *BdFAMA* can compensate for *BdMUTE* in SC recruitment suggested that, in *Brachypodium*, endogenous *BdFAMA* is also poised to drive GMC to GC fate transitions when *BdMUTE* is missing. If this hypothesis were correct, then a double mutant of *bdmute; bdfama* would resemble the single *MUTE* mutants of maize and rice.

We created double mutants using the same *BdFAMA* CRISPR guide and Cas9 vector that we previously used to generate *bdfama*. The regenerants (28 total from 2 independent transformations) were very sick and rarely grew larger than a few millimeters in length before dying and, in many cases, this precluded our ability to collect enough tissue to both image and genotype individuals. While all T0 regenerants generated were lethal, there were two classes of mutant severity. Similar to the maize *bzu2/zmmute*, the severe class of regenerants failed to produce any stomata or recruit SCs, and instead, small-celled GMCs arrested after a few rounds of division (Fig. 4A). Less-severe mutants were able to produce a handful of SC-less stomatal complexes, complete with pore, but the majority of complexes arrested as small cells or paired, undifferentiated, aborted GCs, like those found in *bdmute* (Fig. 4B). We were able to genotype a regenerant from the less severe group, and it was heteroallelic for a 1bp and a 6bp deletion in the second exon of *BdFAMA* (*bdmute*; *bdfama-*Δ*1/*Δ*6)*. The increase in number of aborted GCs in *bdmute; bdfama* mild lines and the complete absence of them in the severe lines suggest that aborted GCs in *bdmute* had, at one time, expressed BdFAMA, but upon longitudinal division, lost expression and were unable to complete differentiation.

**Figure 4.**
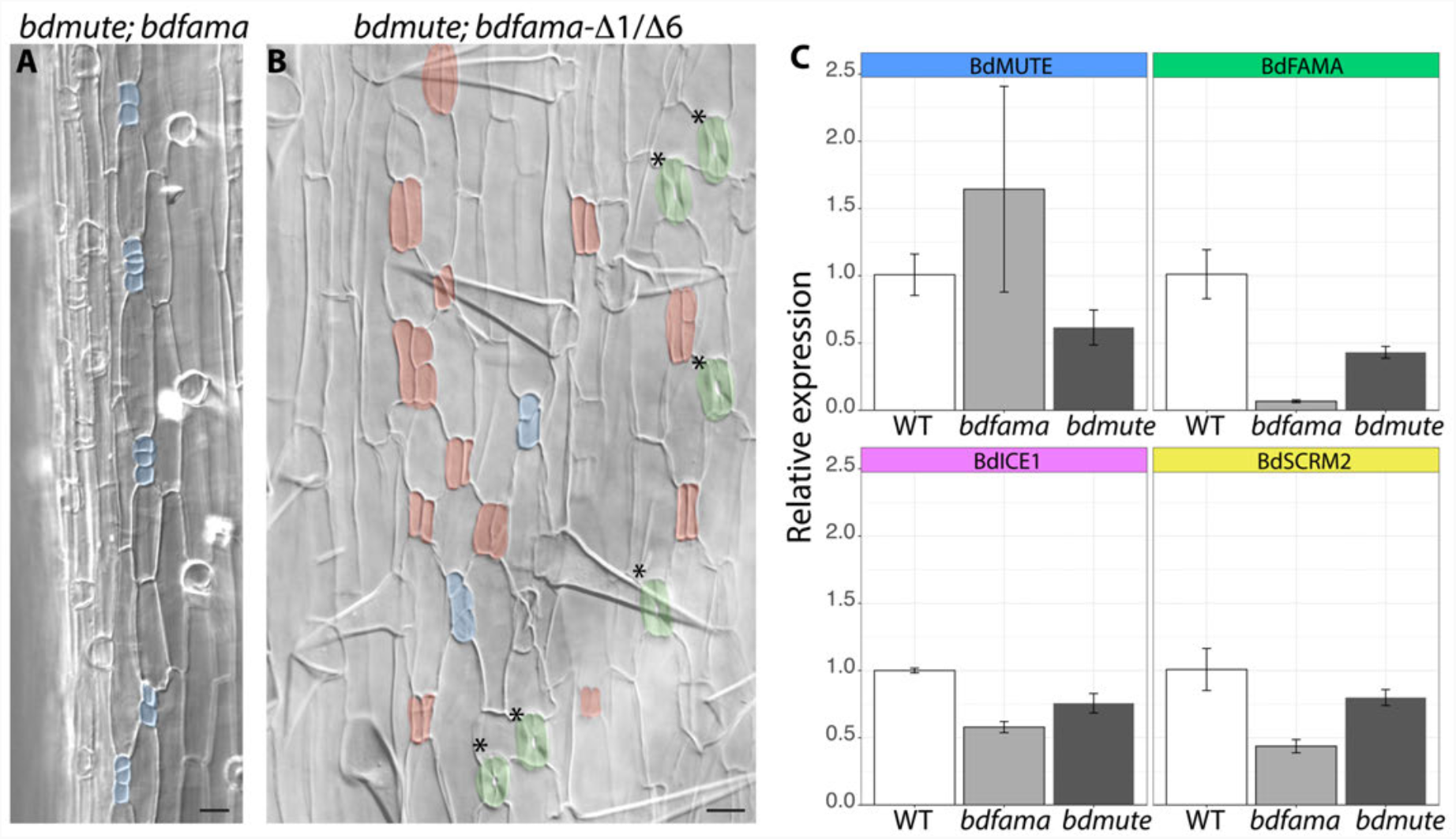
Genetic evidence for differential redundancy of MUTE and FAMA among the grasses. **(A-B)** DIC images of cleared abaxial tissue of *bdmute; bdfama* T0 regenerants. **(A)** Severe *bdmute; bdfama* regenerant with no correctly specified guard cells (GCs) and groups of small cells (blue false color) resembling the stomatal arrest state of rice of maize *mute* mutants **(B)** Milder phenotype of sequence-confirmed heteroallelic *bdmute*; *bdfama-* Δ*1/*Δ*6* with mature stomata lacking subsidiary cells (green false color), aborted GCs (red false color), and arrested small cells (blue false color) **(C)** Relative expression from qRT-PCR of stomatal genes in the leaf division zone of wild type, *bdmute*, and *bdfama*. Expression values were normalized to control gene *BdUBC18* and are relative to expression in wild-type plants. Asterisks indicate significant difference compared to wild-type expression.

### Regulatory relationships within stomatal bHLHs have diverged within the grasses

The genetic and transcriptional data from *Arabidopsis* support a model in which AtMUTE is required for *AtFAMA* expression (Pillitteri *et al*., 2007; Han *et al*., 2018). In maize and rice, *FAMA* expression is reduced significantly in *mute* backgrounds (Wu *et al*., 2019; Wang *et al*., 2019) and suggests that, like in *Arabidopsis*, there is *MUTE-*dependent *FAMA* expression, but our observations of the BdFAMA reporters and phenotypes suggest that there may be a different regulatory strategy in *Brachypodium*. To further define the relationships between *BdMUTE* and *BdFAMA* and their implications for the stomatal lineage, we used qRT-PCR to measure expression levels of stomatal bHLHs in *bdmute, bdfama-*Δ*2* and wildtype (Fig. 4C). *BdFAMA* is readily detectable in *bdmute*, but at a lower level when compared to wildtype. This reduction is consistent with previous findings that 40% of stomatal complexes in *bdmute* fail to differentiate and lack BdFAMA reporter expression (Raissig *et al*., 2017). The *bdmute* allele we used in this study (*bdmute-1/sid*) has a 5bp deletion at position 523-527 in the first exon, and our primers amplify the region between 157-219, so the detection of BdMUTE expression in the *bdmute* background is likely from a partial transcript. The *bdmute-1* allele has been considered a complete loss of function, however, because its mutant phenotype is indistinguishable from CRISPR/Cas9-generated null lines (Raissig *et al*., 2017). *BdSCRM2* and *BdICE1* expression levels are lower in both *bdmute* and *bdfama* backgrounds, but the reduction is more pronounced in the *bdfama* background, which likely corresponds to the reduction of stomatal lineage cells due to failure of stomatal complexes to mature. Remarkably, *BdMUTE* expression increased in *bdfama* compared to wildtype. Together, these data support a model wherein reciprocal regulation between BdFAMA and BdMUTE is required for proper developmental progression, where BdMUTE promotes expression of *BdFAMA* and BdFAMA restrains *BdMUTE*. Although *BdFAMA* levels were reduced in *bdmute*, our ability to detect BdFAMA at all points to a MUTE-independent activation of *BdFAMA* as a reason for BdFAMA’s compensatory abilities, while the MUTE-dependent activation results in lethality in *Arabidopsis*, rice, and maize *mute* mutants.

### *BdFAMA* can rescue GC differentiation but incompletely enforces terminal fate in *Arabidopsis*

FAMA is able to promote GC differentiation in rice, maize, and *Brachypodium*, but it appears that, even within the grasses, the regulation of BdFAMA and BdMUTE has evolved in distinct ways. Across the monocot/dicot divide, key domains of *FAMA* are well-conserved (Fig. S9), and FAMA is similarly required for GC differentiation, but it is additionally required for cell cycle control and terminal fate enforcement in *Arabidopsis*. Given this difference, we asked whether BdFAMA is able to fully or partially rescue cell fate and cell cycle defects in *atfama*. To ascertain degree of rescue, we approached phenotypic analysis with the following criteria: to what extent is BdFAMA able to (1) induce GC fate, (2) prevent excessive GMC divisions, and (3) enforce terminal GC fate? We created lines expressing *AtFAMApro::YFP-BdFAMA* in a heterozygous *atfama/+* background, and genotyping of individual T2 plants indicated that we could obtain living *atfama* homozygous plants, suggesting that BdFAMA can rescue the GC fate defect in *atfama*. These *AtFAMApro::YFP-BdFAMA; atfama* plants had numerous normal-looking stomata, and overall resembled wildtype *Arabidopsis* in stomatal density and patterning (Fig. 5A).

**Figure 5.**
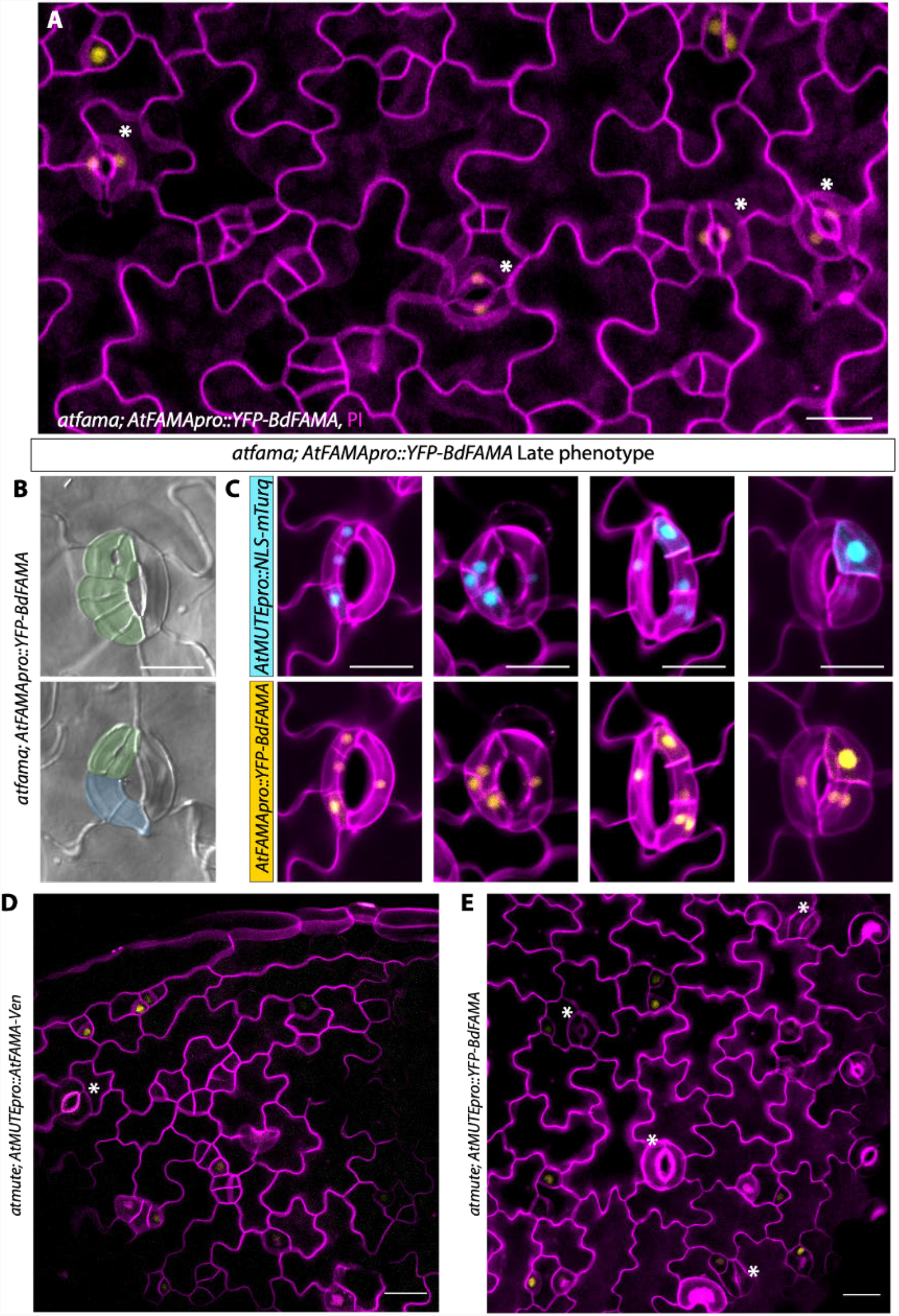
BdFAMA can rescue Arabidopsis guard cell production, but not is not able to fully enforce terminal fate. **(A)** Confocal image of *atfama; AtFAMApro::YFP-BdFAMA* at 5days post germination (dpg) with white asterisks indicating mature stomatal complexes **(B)** DIC images of abaxial cleared tissue from *atfama; AtFAMApro::YFP-BdFAMA* 16 dpg; pseudocolor denotes guard cells (GCs; green) and guard mother cells (GMCs; blue). **(C)** Confocal images of reprogrammed guard cells forming stomata within stomata in late stage (12-16 dpg) abaxial cotyledons from *atfama; AtFAMApro::YFP-BdFAMA; AtMUTEpro::NLS-mTurq*. mTurq (blue) and YFP (yellow) channels shown individually with cell wall stain (magenta). **(D-E)** Confocal images of *atmute; AtMUTEpro::AtFAMA-Ven* and *atmute; AtMUTEpro::YFP-BdFAMA* from 4dpg abaxial cotyledons. **(D)** AtFAMA is able to rescue *atmute*. **(E)** BdFAMA is able to rescue *atmute*. White asterisk indicates stoma. Cell outlines (magenta) visualized by Propidum Iodide (PI) staining. Scale bar=20 μm. **See also Figure S10**

We did observe occasional clusters of small epidermal cells in the rescued lines, but they did not resemble the over-proliferating GC ‘tumors’ typical of *atfama* mutants (Ohashi-Ito *et al*., 2006). To get a more detailed picture of the capacity of BdFAMA to substitute for AtFAMA, we introduced an AtMUTE transcriptional reporter (*AtMUTEpro::NLS-mTurq*) into *atfama; AtFAMApro::YFP-BdFAMA* (Fig. S10A). A timecourse at 3, 4 and 5dpg revealed that the small cells in these clusters and arose from cells that expressed low levels of *AtMUTE* (Fig. S10A, 4dpg) but not all of these cells later expressed BdFAMA and gave rise to stomata; instead, AtMUTE-expressing cells divided again and daughter cells often expressed different levels of *AtMUTE* (Fig. S10A, 5dpg). Expression of AtMUTE demarcates GMC fate and commitment to the stomatal lineage (Pillitteri *et al*., 2007), so the continued division of *AtMUTE*-expressing cells could be caused by the imperfect ability of BdFAMA to drive the GMC fate transition in *Arabidopsis*, but does not necessarily imply a defect in cell division enforcement.

AtFAMA interacts with the *Arabidopsis* homologue of the cell cycle inhibitor, RETINOBLASTOMA RELATED (RBR), through an LxCxE binding motif (Matos *et al*., 2014) (Fig. S9), and this interaction confers terminal GC fate through chromatin modifications (Lee *et al*., 2019). Without this FAMA-RBR interaction, GCs lose their identity and can begin to lobe like pavement cells or divide asymmetrically and reinitiate the stomatal lineage to produce a stomata-in-stomata phenotype (Matos *et al*., 2014). Grass FAMAs have an IxCxE motif (Fig. S9), which may result in a less stable or more transient interaction with RBR (Mcgivern *et al*., 2019). To test whether BdFAMA is sufficient to enforce terminal GC fate, we examined the stomata in the epidermis of older cotyledons (14-17 days post germination (dpg)) in *atfama; AtFAMApro::YFP-BdFAMA* lines and found stomatal complexes that failed to maintain terminal differentiation (Fig. 5B). The BdFAMA-induced phenotype resembled the ‘stomata in stomata’ phenotype produced when the *AtFAMA* LxCxE motif is disrupted, but there are some differences. Notably, GC lobing was less prominent and a single reprogrammed GC could produce several adjacent stomata (Fig. 5B). To better characterize the loss of terminal GC fate, we again examined the *AtMUTE* transcriptional reporter in *atfama; AtFAMApro::YFP-BdFAMA* (Fig. 5C). In stomata whose GCs underwent several divisions, all daughter cells expressed both *AtMUTEpro::NLS-mTurq* and *YFP-BdFAMA*, so it appeared that, rather than reprogramming to the earliest developmental state, with the potential to become a pavement cell or commit to stomatal fate, these daughter cells were only reprogrammed to GMC fate. Newly formed GCs expressing YFP-BdFAMA could also be found immediately adjacent to presumed GMCs, indicating a problem with maintaining one cell-spacing. While BdFAMA is able to rescue stomatal formation and prevent *fama* tumors, the reprogramming of stomata points to a deficiency of BdFAMA to properly regulate the enforcement of terminal GC fate.

The discovery that *BdFAMA* could substitute for *BdMUTE* contrasted with older work in *Arabidopsis* that suggested these two proteins had non-overlapping functions based on a failure to identify viable plants when *AtMUTE* and *AtFAMA* were swapped (Macalister & Bergmann, 2011). *AtFAMA* mutants could, however, be partially rescued by expression of a *Physcomitrium patens* gene that was considered the progenitor of *SPCH, MUTE* and *FAMA* (Macalister & Bergmann, 2011). To test whether BdFAMA could rescue *atmute*, we generated *atmute; AtMUTEpro::YFP-BdFAMA*, and as a specificity control, we also created *atmute; AtMUTEpro::AtFAMA-Venus*. Stomata were successfully produced in both rescue lines, and produced healthy, viable plants (Fig. 5D-E). This suggests that *AtMUTE* and *AtFAMA* have a greater degree of functional overlap than previously appreciated, despite sharing little sequence similarity aside from the conserved bHLH domain and SPCH-MUTE-FAMA (SMF) ACT-like domain (Fig. S9). In both rescue lines, there were adjacent small cells that expressed YFP-BdFAMA, and at 10dpg, clusters of single guard cells and stomata could be seen (Fig. S10 B-C). Both lines produced single GCs, which is indicative of premature transdifferentiation to GC fate. These results demonstrate that, despite millions of years of divergence, *BdFAMA* can substitute for AtFAMA in most GC functions, but also revealed the compensatory ability of *FAMA* for *MUTE* is shared in *Arabidopsis* and *Brachypodium*.

### *BdFAMA* is unable to induce the formation of Brassica-specific myrosin idioblast cells

The ability of BdFAMA to substitute for AtFAMA in GC fate may not be entirely surprising; GCs are a deeply-conserved cell type, so it follows that many of the genes required for GC fate and function are also conserved. But does this functional conservation extend if a transcription factor is co-opted into an new role? *Arabidopsis*, like other Brassicas, is armed with a glucosinolate-myrosinase mustard oil defense to protect against microbes and insects (Shirakawa & Hara-Nishimura, 2018). Both GCs and myrosin idioblasts (MIs) accumulate myrosinases for this defense system, and previous work has shown *AtFAMA* is required for specifying MI fate (Li & Sack, 2014). This cell type is not produced in *Brachypodium* (nor grasses in general), so we were uniquely positioned to ask whether BdFAMA’s ability to rescue *atfama* also extended to this phenotype. MI cells and their nuclei can easily be visualized by expression of a cytoplasmic *AtFAMA* transcriptional reporter (Fig. 6A, D), and by a functional *AtFAMA* translational reporter (Fig. 6B, E), respectively. Although we could readily identify signal in MIs of the control reporter lines and when *AtFAMApro::BdFAMA*-*YFP* was expressed in a wildtype or *atfama/+* background (Fig. S10F), we were unable to detect signal when *AtFAMApro::BdFAMA*-*YFP* was in an *atfama* homozygous background, suggesting that BdFAMA is incapable of rescuing MI fate (Fig. 6C, F). We also visualized mature MI cells using coomassie brilliant blue (CBB) staining and, again, were able to see the characteristic horned shape of mature myrosin cells in the control line, but failed to identify them in the *atfama; AtFAMApro::BdFAMA*-*YFP* lines (Fig. S10D-E).

**Figure 6.**
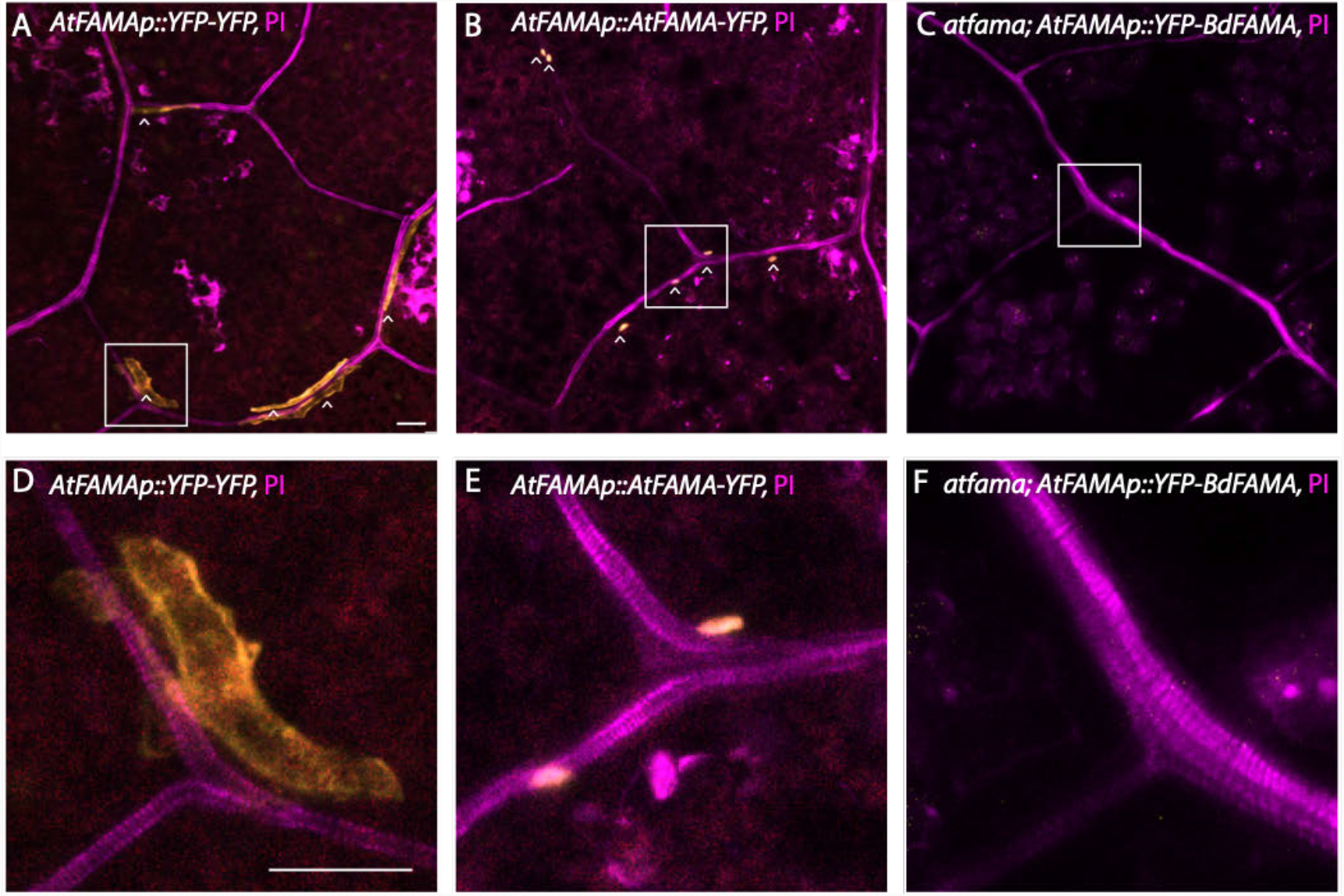
BdFAMA is not able to rescue myrosin cell differentiation in *atfama*. **(A)** Developing myrosin idioblasts (MIs, white carats) expressing transcriptional marker *AtFAMApro::YFP-YFP* (yellow). **(B)** Developing MIs (white carats) showing nuclear expression of translational marker *AtFAMApro::AtFAMA-YFP*. **(C)** *atfama; AtFAMApro::YFP-BdFAMA* fails to rescue MI development and lacks expression of *YFP-BdFAMA*. **(D-F)** Zoomed in images of white box in **(A), (B)**, and **(C)**, respectively. All confocal images are taken from first rosette true leaves 10-15 dpg with Propidum Iodide (PI) cell wall stain. Scale bar=20 μm. **See also Figure S10**

## DISCUSSION

In this study, we found that *BdFAMA* is necessary and sufficient for specifying GC fate in *Brachypodium*. Loss of function mutants fail to differentiate their GCs, form undifferentiated four-celled complexes, and die as a result. Much of what we already knew about FAMA’s role in stomatal differentiation from work in rice and *Arabidopsis* held true in *Brachypodium*, but distinct differences emerged from extending our characterization to encompass FAMA’s interplay with its paralog, MUTE, in these species.

We have phenotypic points of comparison in rice, with *osfama-1*, but lack of extensive analysis of transcriptional and translational reporters and the inconsistency of *fama* phenotypes between rice studies (Liu *et al*., 2009; Wu *et al*., 2019) has precluded the ability to dig deeper into interesting questions; namely, how does a group of well-conserved transcription factors coordinate to produce the patterning and morphology so unique to the grasses? From our expression analysis, we found that BdFAMA’s temporal window begins earlier than expected given its terminal role and what is known from *Arabidopsis*, but, because grasses lack a comparable self-renewing meristemoid stage, the timing is still consistent with expression initiation in GMCs. However, BdFAMA’s maintained expression in mature complexes remains puzzling; does it serve a functional purpose? Is this the case in other grasses? The model in *Arabidopsis* is that the AtFAMA/RBR complex ensures terminal GC fate through deposition of repressive chromatin modifications on stomatal lineage genes such as SPCH and MUTE (Matos *et al*., 2014). Perhaps continued production of BdFAMA in mature stomatal complexes is the strategy grasses use to maintain terminal cell fate, rather than through the repressive chromatin modifications mediated through RBR.

*BdFAMA* is not the only bHLH in *Brachypodium* that has a puzzlingly long expression window for its presumed function: *bdscrm2* has a late arrest phenotype that phenocopies that of *bdfama* and suggests they are heterodimer binding partners, but it is expressed throughout the stomatal lineage, and *bdice1* has an early-stage arrest phenotype similar to the *bdspch1; bdspch2* double mutant, suggesting *BdICE1* forms a heterodimer with *BdSPCHs* (Raissig *et al*., 2016). This supports a strategy where specific heterodimer pairs perform distinct functions, and preferential binding may act as a mechanism for maintaining functional specificity in a lineage with spatiotemporal overlap. This strategy for achieving many functional outcomes from a core set of bHLH TFs has been seen across development, from maize anthocyanin biosynthesis to *Drosophila* neuronal development (Lee et al. 2021, Kong et al. 2021, Powell *et al*. 2008) and recent evidence from *Arabidopsis* hints that the ACT-like domain present in stomatal bHLHs may contribute to binding selectivity (Seo *et al*. 2022). However, binding specificity alone cannot explain how functional specificity is maintained as the interaction data from our CoIP LC-MS/MS assay show all combinations between stomatal group Ia and group IIIb bHLHs are possible *in vivo*.

In a TF family with shared expression timing, heterodimer partners, domains, and the need to coordinate between regulatory and cooperative roles, an additional layer of regulation becomes important. We show that lacking even one functional copy of BdFAMA in the *bdmute* background can cause lethality and mimic the phenotype of *bzu2/zmmute* and *osmute*, and is evidence that *BdFAMA* is indeed able to drive the cell fate transition from GMC to GC in the absence of *BdMUTE* in a dosage-dependent manner. We also observed that severe *bdmute; bdfama* mutant GMCs failed to divide longitudinally to specify GCs, so the aborted GCs seen in the *bdmute* mutant likely expressed BdFAMA earlier in development to drive the GC division, but lost expression upon dividing; providing further evidence of dosage dependence, where late expression of BdFAMA is sensitive to BdFAMA dosage and may require stabilization from BdMUTE. Additionally, while *BdSCRM2* has a late-stage role, it is still able to partially complement *bdice1* to initiate stomatal formation when broadly overexpressed (Raissig *et al*., 2016). Together, these data suggest gene dosage acts as another important layer of regulation in grass stomatal development.

Our genetic comparisons gave us insight, not only into the interplay of key bHLHs of the stomatal lineage, but into the recruitment of grass SCs. The current model for SC recruitment posits that after production of MUTE in GMCs, the TF moves to the neighboring cell file to induce asymmetric division of the SMCs. The demonstrated rescue of *bdmute*’s SC recruitment defect by BdFAMA, however, suggests protein mobility is not an absolute requirement to induce SMC fate. BdFAMA is not mobile, as we did not detect it in any neighboring cell files, so there may instead be mobile intermediaries that BdFAMA is able to activate to prime SMC fate. However, in *bdmute; BdFAMApro::YFP-BdFAMA*, many stomatal complexes fail to properly generate a controlled asymmetric SMC division and instead create abnormal SCs or lobed neighboring cells. This may be due to the inability of *BdFAMA* to substitute for a subset of BdMUTE-specific partners or reflect binding site specificity, or may suggest that movement of the TF to the neighboring cell file is required for consistently oriented polarization of the SMC.

These data reveal a distinction between the ability to induce SMC fate and orient the SMC division. Recent work in *Brachypodium* and maize have identified polarity factors involved in SMC division that support this distinction between fate and division control (Cartwright *et al*., 2009; Wang *et al*., 2019; Zhang *et al*., 2022). In both *Brachypodium* and maize, *Pangloss1* (*PAN1*) polarizes to the SMC/GMC interface and coordinates the nuclear migration and asymmetric SMC division; without it, abnormal SCs and incomplete recruitments occur (Cartwright *et al*., 2009; Zhang *et al*., 2022). An additional polar protein, *BdPOLAR*, was found to oppose the SMC/GMC domain of *BdPAN1* in *Brachypodium* that had an additive effect on SMC division orientation control with *BdPAN1* (Zhang *et al*., 2022). BdPOLAR’s expression in the SMC is dependent on *BdMUTE*, while the expression and polarization of BdPAN1 still occurs in the absence of *BdMUTE*, which explains why we are able to see some successful SC recruitment in *bdmute* lines expressing YFP-BdFAMA. This is also evidence that these two polarity factors have differential dependence on the priming of SMC identity. In the context of our study, BdFAMA is able to compensate for priming SMC identity in the absence of BdMUTE if expressed at the right time, but BdMUTE or its mobility is likely needed for consistent, proper polarization of the SMC asymmetric division.

We have shown that BdFAMA’s compensatory abilities run deeper than initially thought, going so far as to prevent the lethality of *bdmute* that is present in all other characterized plants, including the grasses. We provide evidence that the regulatory relationship between FAMA and MUTE has diverged in *Brachypodium* such that *BdFAMA* is expressed independent of *BdMUTE* and is able to drive the GMC to GC fate transition, but the possibility still remains that there are protein domain differences in BdFAMA that aid in its compensation. In addition to the *how*, the *why* of BdFAMA’s extracurricular role in the GMC to GC fate transition and SC recruitment is intriguing, and presents an opportunity to explore whether this regulatory divergence emerged from domestication events or phylogeny-driven speciation through comparisons of translational and transcriptional reporters in maize and rice. In analyzing the mutant phenotypes of *MUTE* and *FAMA*, it also became apparent that clearer comparisons are needed for developmental stages in the grasses vs. *Arabidopsis*. Grasses do not have a self-renewing meristemoid cell type, but from the *mute* mutant phenotype in maize and rice, and *bdmute; bdfama* double mutant in *Brachypodium*, there does seem to be a pre-GMC stage that requires either *MUTE* or *BdFAMA* to advance. Thus, grass ‘GMCs’ may transition through several distinct developmental stages that may be comparable to *Arabidopsis* development, but inconsistent nomenclature across species obfuscates direct phenotypic comparisons.

This research shows that even one of the most conserved bHLH family members has specialized roles and regulation in its evolution to produce grass stomata. Grasses don’t have the same requirement of dynamic cell cycle control in the stomatal lineage necessary in *Arabidopsis*, and *BdFAMA* seems to have greater stability in mature complexes. Yet, despite millions of years of divergence, *BdFAMA* still seems to retain its cell cycle control, even though the *Brachypodium* loss-of-function phenotype suggests it is not a required role for the lineage. With BdFAMA expression occurring so early in the lineage, it stands to reason that having a strong cell cycle inhibitory role may prove detrimental if stochastic levels of BdFAMA can cause aberrant GMC division failure and reduce stomatal numbers. The grass leaf is also set up with much stronger developmental zoning than in *Arabidopsis*, and the grass stomatal lineage does not have dividing stem cell-like populations that demand high regulatory plasticity, so cell division control along the leaf may defer to a more global regulatory signal.

The MI cells are another example of functional divergence between *Arabidopsis* (Brassica) and grasses. Here the role of FAMA is different; in *Arabidopsis*, it is expressed in small ground meristem cells and is required, along with AtICE1 or AtSCRM2, for myrosin cell development (Li & Sack, 2014; Shirakawa *et al*., 2016). The requirement of these three bHLHs for myrosin cell differentiation indicate the presence of a shared transcription factor network between myrosin cells and GCs, and yet *BdFAMA* was unable to rescue the formation of myrosin cells and we failed to detect expression in the absence of AtFAMA. Guard cell vacuoles in Brassicales plants also contain myrosinases, so it is thought that *FAMA* was first co-opted into this defense system in stomata, and later, during evolution, developed myrosin cells (Shirakawa & Hara-Nishimura, 2018). Myrosin cells represent a synapomorphy in Brassicas and although BdFAMA is unable to participate in their formation, the fact that we were able to observe expression of *AtFAMApro::YFP-BdFAMA* in wildtype and *atfama/+* lines hints at a myrosin-specific regulatory strategy. Specifically, expression of *AtFAMA* in the *Arabidopsis* stomatal lineage is MUTE-dependent and FAMA-independent (Ohashi-Ito *et al*., 2006; Han *et al*., 2018), however, *AtMUTE* expression is not detected in myrosin cells (Li & Sack, 2014), so there is a difference in the regulatory relationship between MUTE and FAMA within the *Arabidopsis* epidermis and inner leaf tissue. We propose that, in myrosin cells, AtFAMA is required for its own expression, as we failed to detect expression of *YFP-BdFAMA* in the absence of AtFAMA, but could see expression in wildtype and *atfama/+* backgrounds. Domain swaps between *AtFAMA* and *BdFAMA* will help uncover structural features responsible for the differential capabilities, and comparative transcriptomics of *atfama* mutants rescued by AtFAMA or BdFAMA may be useful to determine what genes are required to make MIs.

This study demonstrated the functional conservation of *BdFAMA* in driving GC fate across the monocot/dicot divide, but the difference seen in *MUTE* mutant phenotypes and differential regulation of *FAMA* across the grasses brings to light that our ability to understand the role of one gene hinged on understanding the system as a whole. We show that with thorough investigation of transcriptional and translational markers, in addition to leveraging genetic manipulations within and across species, it is possible to get a more complete picture of the players in pathways like the stomatal lineage. Furthermore, the existence of reporter lines for each bHLH transcription factor in the *Brachypodium* stomatal lineage is what enabled our pursuit of *in vivo* proteomics that produced interaction data useful for identifying novel candidates involved in stomatal development and regulation. A systematic understanding of plant systems and their mechanistic underpinnings is needed to improve crops, and this relies on our ability as a field to produce high quality proteomics resources and careful genetic evaluation using technologies like CRISPR/Cas9 gene editing. As a temperate grass model, *Brachypodium distachyon* can provide a framework from which future studies on upstream environmental-originating signaling and downstream stomatal effectors can stem, and can contribute to improvements in agriculturally significant temperate grasses like wheat and barley.

## MATERIALS AND METHODS

### Plant Material and Growth Conditions

The *Brachypodium* line, Bd21-3, was used as the wildtype background in all experiments (Vogel & Hill, 2008). Plants were generally grown as specified in Raissig *et al*. 2016 and 2017, briefly, seeds were sterilized for 15 minutes in a solution of 20% bleach and 0.1% Tween, and stratified on 1/2 strength MS agar plates (Cassion Labs, 1% Agar, pH 5.7) for at least 2 days at 4°C in dark before being transferred to 22°C chamber with a 16-hour light/8-hour dark cycle (110 μmol m^−2^ s^−1^). Plants on soil were grown in 4×4-inch pots in a greenhouse with a 20-hour light/4-hour dark cycle (250-300 μmol m^−2^ s^−1^; day temperature = 28°C, night temperature = 18°C), with the exception of lines *sid/bdmute-1; BdMUTEpro::YFP-BdFAMA* and *wildtype; BdMUTEpro::YFP-BdFAMA*, which were grown on soil in a Percival growth room at 26°C with a 16-hour light/8-hour dark cycle (250 μmol m^−2^ s^−1^). The *bdmute* mutant used in this study corresponds to the sid/bdmute-1 allele from Raissig et al. 2017. Columbia (Col-0) was used as the wildtype background for all *Arabidopsis* experiments, except for *atmute/+*, which is the segregating G→A mutant allele from an EMS mutagenized population used in Pilliteri et al. 2007. *atfama/+* is the segregating T-DNA insertion line Salk_100073. *Arabidopsis* seeds were surface sterilized and grown on 1/2 MS plates, as described above. *Arabidopsis* plants grown on soil were grown in 22°C chambers as above, then transferred to light racks with 16-hour light/8-hour dark cycle (110 μmol m^−2^ s^−1^).

### Generation of DNA Constructs

The following constructs for use in *Brachypodium* were generated in this study: *BdFAMApro::YFP-BdFAMA, bdfama Cas9-g4, Ubipro::YFP-BdFAMA, Ubipro::BdFAMA, BdFAMApro::3xYFP, BdMUTEpro::YFP-BdFAMA*. To create *BdFAMApro:YFP-BdFAMA*, ∼4.7 kb upstream sequence of the *BdFAMA* gene (Bradi2g22810; primers primMXA7-FWD and primMXA8-REV) was cloned into pIPKb001t (Raissig et al., 2016). Separately, the *BdFAMA* genomic sequence (primers primMXA5-FWD and primMXA6-REV) was cloned into pENTR-D with a poly-alanine linker (annealed primers Ala_linker-F and Ala_linker-R) and an AscI-flanked *Citrine* YFP inserted 3’ of the gene by AscI digest. Site-directed mutagenesis with primers primMXA20 and primMXA21 was used to eliminate a NotI recognition site in the second exon that interfered with the cloning of BdFAMA. Finally, the entry clone was recombined into the pIPKb001t vector. *BdFAMApro::3xYFPnls* was created as above, except with the *BdFAMA* genomic sequence substituted with 3xYFPnls pENTR. *Ubipro::YFP-BdFAMA* and *Ubipro::BdFAMA* were made using pIPK002, the monocot overexpression Gateway vector, as a backbone. *BdMUTEpro::YFP-BdFAMA* was created by recombining the pENTR_YFP-BdFAMA with the destination vector, pIPKb001_BdMUTEpro, used in Raissig et al. 2017. We created the CRISPR constructs using the vectors pH-Ubi-cas9-7 and pOs-sgRNA (vectors and protocol described in Miao et al., 2013). The online server tool, CRISPR-P, was used to identify candidate spacer sequences (Lei et al., 2014). Spacers were generated by annealing oligo duplexes primMXA3 and primMXA4 for *BdFAMA* CRISPR_sgRNA_4, which, along with pH-Ubi-cas9-7 were transformed into Bd21-3 or *bdmute* to generate *bdfama-*Δ*2* and *bdmute*; *bdfama*, respectively. Primers primMXA22 and 24 were used to genotype the area flanking the predicted cut site. All primers are listed in Table S1.

The following constructs were generated for use in Arabidopsis lines: *AtFAMApro::YFP-BdFAMA, AtFAMApro::AtFAMA-YFP, AtMUTEpro::YFP-BdFAMA, AtMUTEpro::NLS-mTurq*, and *AtFAMApro::NLS-mTurq*. Constructs for *Arabidopsis* transformation were created using Gateway LR recombination. R4pGWB501, pDONR_L1_AtMUTEpro_R4 or pJET_L4_AtFAMApro_R1 (∼2.4kb upstream of AtFAMA start codon), and pENTR_YFP-BdFAMA were recombined to form constructs *AtMUTEpro::YFP-BdFAMA and AtFAMApro::YFP-BdFAMA*, respectively. *AtFAMApro::AtFAMA-YFP* was created by recombining R4pGWB640, pJET_L4_AtFAMApro_R1, and pENTR_L1_gAtFAMA_L2 (cloned from gDNA with primers gFAMA-FWD and gFAMA-REV). *AtMUTEpro::AtFAMA-Ven* was acquired from Margot Smit, personal communication. The Greengate cloning system (Lampropoulos *et al*., 2013) was used to make *Arabidopsis* transcriptional markers, *AtMUTEpro::NLS-mTurq, AtFAMApro::NLS-mTurq* with sulf resistance in the *pGreen0229* backbone.

We created the CRISPR constructs using the vectors pH-Ubi-cas9-7 and pOs-sgRNA (vectors and protocol described in Miao et al., 2013). The online server tool, CRISPR-P, was used to identify candidate spacer sequences (Lei et al., 2014). Spacers were generated by annealing oligo duplexes primMXA3 and primMXA4 for *BdFAMA* CRISPR_sgRNA_4, which, along with pH-Ubi-cas9-7 were transformed into Bd21-3 or *bdmute* to generate *bdfama-*Δ*2* and *bdmute*; *bdfama*, respectively. Primers primMXA22 and 24 were used to genotype the area flanking the predicted cut site. All primers are listed in Table S1.

### Generation of transgenic plant lines

Transgenic lines were created from *Brachypodium* calli derived from Bd21-3 and *bdmute* parental plants, transformed with *Agrobacterium tumefaciens* strain AGL1, selected for hygromycin resistance, and regenerated according to standard *Brachypodium* tissue culture protocols (http://jgi.doe.gov/our-science/scienceprograms/plant-genomics/brachypodium/); regenerated plants from calli are referred to as T0 plants, and their immediate progeny as T1. Leaves from T0 regenerants bearing a fluorescent reporter were stained with Propidium Iodide (PI) (1:100 dilution of 1 mg/ml stock Molecular Probes; Invitrogen detection technologies) and visually inspected using confocal microscopy for expression of the reporter as confirmation of successful transformation. The floral dip protocol with GV301 *Agrobacterium* strain was used to transform *Arabidopsis* plants of Col-0, *atfama/+*, or *atmute/+* backgrounds (Clough et al 1998). *atmute; AtMUTEpro::AtFAMA-Ven* was made by crossing *AtMUTEpro::AtFAMA-Ven* (line from Margot Smit, pers. comm) into the *atmute/+* mutant background using standard procedures (Weigel & Glazebrook, 2006). All lines are listed in Table S2.

### Microscopy and Phenotypic Analysis

All *Brachypodium* and *Arabidopsis* confocal imaging was done using a Leica SP5 or Stellaris confocal microscope, as described in Raissig et al 2016. In sum, the youngest emerging leaves were carefully excised with forceps from the surrounding sheath and stained in PI (1:100 dilution of 1 mg/ml stock Molecular Probes; Invitrogen detection technologies) to visualize cell walls, and were mounted in water to image the abaxial leaf surface. For quantification of the nuclear signal intensity of *BdMUTEpro::YFP-BdFAMA*, the same laser power and settings were used across all samples. Reporter expression levels and cell size change depending on stomatal developmental stage, but within early developmental stages (after asymmetric division and before SC recruitment), cell shape is relatively consistent, so we standardized the quantification by only measuring the fluorescent intensity in the nuclei of cells whose length and width fell between 4 and 6 μm. FIJI was then used to SUM-project images and measure the integrated density of nuclear signal. To observe fluorescent signal in myrosin cells, young true leaves were collected and stained in PI, as above, for 15 minutes, and firmly pressed when mounted with a glass coverslip, abaxial side up, in water.

For all DIC imaging, either a Leica DM6 B microscope or a Leica DMi8 inverted scope were used. To visualize tissue for DIC imaging and remove chlorophyll, *Brachypodium* leaves were collected and placed in 7:1 ethanol:acetic acid solution overnight, at minimum, rinsed with water, and mounted in Hoyer’s medium. The same was done with *Arabidopsis* tissue, except tissue was left in solution for at least 2 hours before rinsing and mounting to clear. Quantification of cell numbers from cleared tissue were done with images captured on the inverted Leica DMi8 inverted scope using a 0.29mm^2^ field of view; in general, 5 fields of view per leaf and 6 leaves from 6 individuals were used. To visualize the leaf morphology of regenerants in BdFAMA overexpression lines, regenerants still attached to calli were visually inspected using a dissecting scope.

### Coomassie Brilliant Blue staining of myrosin cells

To visualize mature myrosin cells in *Arabidopsis*, true leaves of 10-15 day old plants were stained as described in Ueda *et al*., 2006: leaves were boiled for 3 minutes in Coomassie Brilliant Blue (CBB) solution (45% methanol, 10% acetic acid, and 0.25% CBB R250), and left to destain for 2-4 hours in Hoyer’s medium. After destaining, tissue was mounted on glass slides with 60% glycerol and the abaxial surface examined with DIC microscopy for myrosin cells.

### DNA extraction and qRT-PCR

When DNA was extracted for genotyping using the CTAB DNA extraction protocol (Allen et al. 2006), tissue was harvested and snap-frozen with glass beads in liquid nitrogen, and ground using a Spex Certiprep Geno Grinder 2000. Alternatively, DNA was extracted using the Phire^™^ Plant Direct PCR Master Mix Kit (ThermoScientific), following the manufacturer’s instructions, using ∼2-5mm diameter tissue sample obtained from a young leaf suspended in 50uL of the provided dilution buffer and either ground with the tip of a pipette or boiled for 10 minutes. Sanger sequencing was used following PCR for genotyping; genotyping primers can be found in Table S1. We extracted RNA for use in qRT-PCR from wildtype, *bdmute*, and *bdfama* leaves from the division zone (3 individuals per replicate, except for *bdfama*, which had 2 individuals per replicate, 3 replicates; the first leaf was removed and ∼4mm of the base of emerging leaves, starting from the shoot apical meristem, were sampled) using the RNeasy Mini kit (Qiagen) following the manufacturer’s instructions, including optional DNase treatment. To produce single stranded cDNA, we used the iScript™ cDNA Synthesis Kit (Bio-Rad) and quantified the cDNA using a Nanodrop (ThermoScientific, NanoDrop 1^c^). For qRT-PCR we used the SsoAdvanced™ Universal SYBR^®^ Green Supermix (Bio-Rad) and a CFX96™ Real Time System (Bio-Rad) running a standard program (initially 95°C for 30sec, then 95°C for 10sec, 60°C for 30sec, 40 cycles, followed by a melting curve). All qRT-PCR primers were designed using QuantPrime (www.quantprime.de); primer sequences can be found in Table S1.

### Protein Alignments

Protein alignment of *BdFAMA* (Bradi2g22810), *BdMUTE* (Bradi1g18400), *AtFAMA* (AT3G24140), and *AtMUTE* (AT3G06120) were done using the clustal 2.1 multiple sequencing alignment (MSA) tool (https://www.ebi.ac.uk/Tools/msa/). *Brachypodium* sequences were obtained from the *Brachypodium distachyon v3*.*1* in Phytozome 10. Protein domains were identified using the National Center for Biotechnology Information conserved domain search tool (https://www.ncbi.nlm.nih.gov/Structure/cdd/wrpsb.cgi).

### Yeast two hybrid (Y2H) tests

To obtain cDNA suitable for creating Y2H compatible *Brachypodium* genes, Bd21-3 seedlings were grown on 1/2 MS plates and the bottom 5 mm (developmental zone) of young true leaves were harvested, flash frozen and ground for RNA extraction using the RNeasy Plant MiniKit (Quiagen), followed by reverse transcription with SuperScript IV reverse transcriptase (Thermo Fisher) according to the manufacturer’s instructions. Genes with stop codons were amplified using the primers in Table 1 and cloned into pENTR/D-TOPO, followed by LR recombination with gateway compatible pGAD-T7 and pGBK-T7 (see vector maps). For Y2H assays, bait and prey plasmids were transformed into the yeast strain AH109, followed by selection of transformants and testing of pair-wise interactions by growth complementation assays on nutrient selection media as described in the Matchmaker™ GAL4 Two-Hybrid System 3 manual (Clontech). To compensate for auto-activation of some constructs, selective plates containing 10 – 30 mM 3-AT were included in the interaction assays.

### AP-MS

#### Plant lines

*Brachypodium* Bd21-3 wild type and the following published reporter lines in the wild type background were used for AP-MS: BdSPCH2pro::BdSPCH2-CitYFP (Raissig *et al*., 2016), BdMUTEpro::CitYFP-BdMUTE (construct and transformation strategy described in (Raissig *et al*., 2017)), BdSCRM2pro::CitYFP-BdSCRM2 (Raissig *et al*., 2016), ZmUBI1pro::YFP-BdICE1 (Raissig *et al*., 2016), BdMUTEpro::3xYFPnls (Raissig *et al*., 2017). The creation of BdFAMApro::YFP-BdFAMA is described above.

#### Growth and harvesting conditions

For each sample, about 100 seeds were surface sterilized with 1 % bleach, 0.1 % Tween-20 for 10 min, washed thoroughly with water, stratified on ½ MS plates (Caisson labs, 1% Agar, pH 5.7) for 4-6 days at 4°C and transferred to a growth chamber set to long day conditions (16 h/8 h light/dark, 22°C). Starting from the 2^nd^ leave until the 5^th^ or 6^th^ main shoot leaf, all emerging leaves were gently pulled off the plant and the bottom 4 mm containing the developmental zone (as determined by microscopy) were harvested, flash frozen in liquid nitrogen and stored at -80°C until further use. Plants were returned to the chamber to grow additional leaves after each round of harvesting. In total, about 150-220mg of plant material from 600-700 leaves were obtained per sample.

#### Affinity purification

For each line, three biological replicates were used. Frozen plant material was ground to a fine powder in a ball mill, weighed and resuspended in 5 µl/mg extraction buffer (25 mM Tris pH 7.5, 10 mM MgCl2, 0.5 mM EGTA, 75 mM NaCl, 1 % Triton-X-100, 1 mM NaF, 0.5 mM Na3VO4, 15 mM beta-Glycerophosphate, 1 mM DTT, 1 mM PMSF, 1x complete proteasome inhibitor) by rotating at 4°C for 15 min. Cells were ruptured by 5 × 15 sec sonication in an ice bath in Bioruptor UCD-200 (Diagenode) at medium intensity with 2 min breaks on ice. To further break cell walls and digest DNA, 2 µl Lysonase (Millipore) were added and samples were incubated at 4°C for 30 min with endo-over-end rotation. The extracts were cleared by 10 min centrifugation at 12,000 rpm and the supernatant was incubated on a rotor wheel at 4°C for 3 h with 25 µl of GFP-Trap magnetic agarose bead slurry (Chromotek), pre-washed with extraction buffer. The beads were washed tree times with 500 µl wash buffer (extraction buffer with 100 mM NaCl and without proteasome inhibitor), resuspended in 50 µl 2x Leammli buffer, boiled for 5 min at 95°C and frozen at -80°C until further processing. Successful affinity purification was confirmed by western blotting using Rat monoclonal anti-GFP antibody (3H9, Chromotek) and AffiniPure Donkey Anti-Rat IgG-HRP (712-035-153, Jackson Immuno Research Laboratories).

#### Sample prep and mass spectrometry

For MS analysis, purified proteins were briefly separated by SDS-PAGE on Mini PROTEAN TGX gels (BioRad), stained with the Novex colloidal blue staining kit (Invitrogen) and excised and further processed by in-gel Tryptic digestion. Peptides were desalted using C18 ZipTips (Millipore) and resuspended in 0.1% formic acid. LC-MS/MS was performed on a Q-Exactive HF hybrid quadrupole-Orbitrap mass spectrometer (Thermo Fisher) with an Easy LC 1200 UPLC liquid chromatography system (Thermo Fisher) and analytical column ES803 (Thermo Fisher). Peptides were eluted with the following conditions at a flow rate of 300 nl/min: a 100 min gradient from 5 to 28 % solvent B (80 % acetonitrile, 0.1 % formic acid), followed by a 20 min gradient from 28 to 44 % solvent B and a short wash with 90 % solvent B. Precursor scan was from mass-to-charge ratio (m/z) 375 to 1,600 and the top 20 most intense multiply charged precursors were selection for fragmentation with higher-energy collision dissociation (HCD) with normalized collision energy (NCE) 27.

#### Data analysis

Protein identification and label-free quantification (LFQ) were done in MaxQuant (version 1.6.2.6) (Tyanova *et al*., 2016a) using the default settings with the following modifications: LFQ minimum ratio count was set to 1, “Fast LFQ” and “Match between runs” were enabled. Peptides were searched against the *B. distachyon* Bd21-3 protein database (v1.1) obtained from Phytozome containing a total of 47,917 entries (https://phytozome-next.jgi.doe.gov/info/BdistachyonBd21_3_v1_1) plus a list of likely contaminants containing Trypsin, human Keratin, and YFP and against the contaminants list of MaxQuant. Search results were then analyzed in Perseus (version 1.6.2.3) (Tyanova *et al*., 2016b). LFQ intensities were imported and proteins marked as ‘only identified by site’, ‘reverse’, ‘potential contaminant’ and such that were not identified in at least two replicates of one sample group were removed. Data were log2 transformed and missing values imputed from normal distribution (width = 0.3, down shift = 1.8, total matrix mode). Significantly enriched proteins were identified by unpaired 2-sided Students t-tests (modified permutation-based FDR with 250 randomizations for multiple sample correction; S0 = 0.5) comparing the bHLH reporter lines with either of the two control lines. Only proteins that were identified in at least two replicates of the bHLH reporter line were used for the test. False discovery rates (FDRs) were chosen to get a minimum number of ‘false negatives’: 0.015 for BdSPCH2 vs. Bd21-3, 0.025 for BdMUTE/BdICE1/YFP vs. Bd21-3, 0.05 for BdFAMA/BdSCRM2 vs. Bd21-3, 0.075 for BdSPCH2/BdMUTE/BdFAMA/BdSCRM2 vs. YFP and 0.04 for BdICE1 vs. YFP and 0.035 for Bd21-3 vs. YFP. To be considered bHLH interaction candidates, proteins had to meet the following criteria: (1) they had to be identified by at least one MS/MS spectrum in the respective sample group, (2) they had to be significantly enriched vs. at least one of the controls and (3) the fold change in the bHLH samples had to be at least 1.5 times higher than that of the other control or 2 times if the other control was also significantly enriched. Data were also searched with Protein Prospector using the same databases and search parameters described in (Shrestha *et al*., 2022). Search results were compared using a protein and peptide FDR cutoff of 5% and 1% with keep replicates enabled to enable. Peptide counts of candidates are largely in agreement with MaxQuant results.

OrthoFinder (Emms & Kelly, 2019) and the *Brachypodium* gene IDs (circa 2019) were used to identify orthologs of candidate proteins in other plant species. Gene ontology (GO) terms (biological processes) for rice and *Arabidopsis* genes were obtained from uniprot. Additional GO analysis of the *Brachpodium* candidates and their best rice and *Arabidopsis* orthologs was done with AgriGO v2.0 (Tian *et al*., 2017). Likely protein function was deduced from GO term analysis and functional annotation on TAIR (Arabidopsis.org). Hierarchical clustering and principal component analysis (PCA) were done in Perseus. PCA, scatter plots and Venn diagrams were plotted in RStudio (RStudio, 2022). The MS data (raw files and search results) will be made accessible through the ProteomeXchange Consortium (http://proteomecentral.proteomexchange.org) upon publication.

### Statistical Analysis and Plotting

Statistical analysis was performed in R. For phenotypic quantifications, two sample tests (ie. Wildtype line vs mutant line or reporter line A fluorescence vs reporter line B fluorescence) were performed using the Wilcoxon-rank sum test followed by Dunn’s multiple comparisons test. To analyze qRT-PCR data, a Welch’s t test was applied to compare mutant line relative to wildtype levels.

## ACKNOWLEDGEMENTS

We thank members of the lab for creative and critical discussions about genetic and developmental diversity, Dr. Michael Raissig for access to data, reagents and discussions of Brachypodium stomatal regulation, Dr. Margot Smit for Arabidopsis cell fate reporters pre-publication, Dr. Yan Gong for image analysis expertise and Joel Erberich for help with analyzing gene expression. KHM was supported by an NSF graduate research fellowship and by NIHGRI training grant 5T32HG000044 awarded to the Stanford University School of Medicine. DCB is an HHMI investigator.

## REFERENCES

Adrian J, Chang J, Ballenger CE, Bargmann BOR, Alassimone J, Davies KA, Lau OS, Matos JL, Hachez C, Lanctot A, et al. 2015. Transcriptome dynamics of the stomatal lineage: birth, amplification and termination of a self-renewing population. Developmental cell 33: 107.

Bergmann DC, Lukowitz W, Somerville CR. 2004. Stomatal development and pattern controlled by a MAPKK kinase. Science 304: 1494–1497.

Cartwright HN, Humphries JA, Smith LG. 2009. PAN1: A receptor-like protein that promotes polarization of an asymmetric cell division in maize. Science 323: 649–651.

Chater CC, Caine RS, Tomek M, Wallace S, Kamisugi Y, Cuming AC, Lang D, MacAlister CA, Casson S, Bergmann DC, et al. 2016. Origin and function of stomata in the moss Physcomitrella patens. Nature plants 2: 16179.

Davies KA, Bergmann DC. 2014. Functional specialization of stomatal bHLHs through modification of DNA-binding and phosphoregulation potential PLANT BIOLOGY. PNAS 111: 15585–15590.

Emms DM, Kelly S. 2019. OrthoFinder: Phylogenetic orthology inference for comparative genomics. Genome Biology 20: 1–14.

Geisler M, Nadeau J, Sack FD. 2000. Oriented asymmetric divisions that generate the stomatal spacing pattern in Arabidopsis are disrupted by the too many mouths mutation. Plant Cell 12: 2075–2086.

Hachez C, Ohashi-Ito K, Dong J, Bergmann DC. 2011. Differentiation of arabidopsis guard cells: Analysis of the networks incorporating the basic helix-loop-helix transcription factor, FAMA. Plant Physiology 155: 1458–1472.

Han S-KK, Qi X, Sugihara K, Dang JH, Endo TA, Miller KL, Kim ED, Miura T, Torii KU. 2018. MUTE directly orchestrates cell state switch and the single symmetric division to create stomata. Developmental Cell 45: 303-315.e5.

Kanaoka MM, Pillitteri LJ, Fujii H, Yoshida Y, Bogenschutz NL, Takabayashi J, Zhu J-K, Torii KU. 2008. SCREAM/ICE1 and SCREAM2 specify three cell-state transitional steps leading to Arabidopsis stomatal differentiation. The Plant Cell 20: 1775–1785.

Lampropoulos A, Sutikovic Z, Wenzl C, Maegele I, Lohmann JU, Forner J. 2013. GreenGate---a novel, versatile, and efficient cloning system for plant transgenesis. PloS one 8.

Lee E, Lucas JR, Goodrich J, Sack FD. 2014. Arabidopsis guard cell integrity involves the epigenetic stabilization of the FLP and FAMA transcription factor genes. The Plant Journal 78: 566–577.

Lee LR, Wengier DL, Bergmann DC. 2019. Cell-type–specific transcriptome and histone modification dynamics during cellular reprogramming in the Arabidopsis stomatal lineage. Proceedings of the National Academy of Sciences of the United States of America 116: 21914–21924.

Li M, Sack FD. 2014. Myrosin idioblast cell fate and development are regulated by the Arabidopsis transcription factor FAMA, the auxin pathway, and vesicular trafficking. Plant Cell 26: 4053–4066.

Liu T, Ohashi-Ito K, Bergmann DC. 2009. Orthologs of Arabidopsis thaliana stomatal bHLH genes and regulation of stomatal development in grasses. Development (Cambridge, England) 136: 2265–76.

Lopez-Anido CB, Vatén A, Smoot NK, Sharma N, Guo V, Gong Y, Anleu Gil MX, Weimer AK, Bergmann DC. 2021. Single-cell resolution of lineage trajectories in the Arabidopsis stomatal lineage and developing leaf. Developmental Cell 56: 1043-1055.e4.

Macalister CA, Bergmann DC. 2011. Sequence and function of basic helix-loop-helix proteins required for stomatal development in Arabidopsis are deeply conserved in land plants. Evolution and Development 13: 182–192.

MacAlister CA, Ohashi-Ito K, Bergmann DC. 2007. Transcription factor control of asymmetric cell divisions that establish the stomatal lineage. Nature 445: 537–540.

Mair A, Xu SL, Branon TC, Ting AY, Bergmann DC. 2019. Proximity labeling of protein complexes and cell type specific organellar proteomes in Arabidopsis enabled by TurboID. eLife 8.

Matos JL, Lau OS, Hachez C, Cruz-Ramírez A, Scheres B, Bergmann DC. 2014. Irreversible fate commitment in the Arabidopsis stomatal lineage requires a FAMA and RETINOBLASTOMA-RELATED module. eLife 3: e03271.

Mcgivern DR, Findlay KC, Montague NP, Boulton MI. 2019. An intact RBR-binding motif is not required for infectivity of Maize streak virus in cereals, but is required for invasion of mesophyll cells. Journal of General Virology 16: 42.

Negi J, Moriwaki K, Konishi M, Yokoyama R, Nakano T, Kusumi K, Hashimoto-Sugimoto M, Schroeder JI, Nishitani K, Yanagisawa S, et al. 2013. A Dof Transcription Factor, SCAP1, Is Essential for the Development of Functional Stomata in Arabidopsis. Current Biology 23: 479–484.

Ohashi-Ito K, Bergmann DC, Nagano AJ, Shimada T, Kohchi T, Hara-Nishimura I, Oa W, Ohashi-Ito K, Bergmann DC. 2006. Arabidopsis FAMA controls the final proliferation/ differentiation switch during stomatal development. The Plant cell 18: 2493–505.

Ortega A, de Marcos A, Illescas-Miranda J, Mena M, Fenoll C. 2019. The tomato genome encodes SPCH, MUTE, and FAMA candidates that can replace the endogenous functions of their Arabidopsis orthologs. Frontiers in Plant Science 10: 1300.

Pillitteri LJ, Sloan DB, Bogenschutz NL, Torii KU. 2007. Termination of asymmetric cell division and differentiation of stomata. Nature 445: 501–505.

Pires N, Dolan L. 2010. Origin and diversification of basic-helix-loop-helix proteins in plants. Molecular Biology and Evolution 27: 862–874.

Raissig MT, Abrash E, Bettadapur A, Vogel JP, Bergmann DC. 2016. Grasses use an alternatively wired bHLH transcription factor network to establish stomatal identity. Proceedings of the National Academy of Sciences of the United States of America 113: 8326–31.

Raissig MT, Matos JL, Anleu Gil MX, Kornfeld A, Bettadapur A, Abrash E, Allison HR, Badgley G, Vogel JP, Berry JA, et al. 2017. Mobile MUTE specifies subsidiary cells to build physiologically improved grass stomata. Science (New York, N.Y.) 355: 1215–1218.

RStudio T. 2022. RStudio Team Overview -RStudio Documentation. PBC: Boston, MA.

Shirakawa M, Hara-Nishimura I. 2018. Specialized Vacuoles of Myrosin Cells: Chemical Defense Strategy in Brassicales Plants. Plant and Cell Physiology 59: 1309–1316.

Shirakawa M, Ueda H, Shimada T, Hara-Nishimura I. 2016. Myrosin cells are differentiated directly from ground meristem cells and are developmentally independent of the vasculature in Arabidopsis leaves. Plant signaling & behavior 11: e1150403.

Shrestha R, Reyes A V., Baker PR, Wang ZY, Chalkley RJ, Xu SL. 2022. 15N Metabolic Labeling Quantification Workflow in Arabidopsis Using Protein Prospector. Frontiers in Plant Science 13: 46.

Tian T, Liu Y, Yan H, You Q, Yi X, Du Z, Xu W, Su Z. 2017. agriGO v2.0: a GO analysis toolkit for the agricultural community, 2017 update. Nucleic acids research 45: W122–W129.

Tyanova S, Temu T, Cox J. 2016a. The MaxQuant computational platform for mass spectrometry-based shotgun proteomics. Nature Protocols 2016 11:12 11: 2301–2319.

Tyanova S, Temu T, Sinitcyn P, Carlson A, Hein MY, Geiger T, Mann M, Cox J. 2016b. The Perseus computational platform for comprehensive analysis of (prote)omics data. Nature Methods 2016 13:9 13: 731–740.

Vizcaíno JA, Côté RG, Csordas A, Dianes JA, Fabregat A, Foster JM, Griss J, Alpi E, Birim M, Contell J, et al. 2013. The PRoteomics IDEntifications (PRIDE) database and associated tools: status in 2013. Nucleic acids research 41.

Vogel J, Hill T. 2008. High-efficiency Agrobacterium-mediated transformation of Brachypodium distachyon inbred line Bd21-3. Plant cell reports 27: 471–478.

Wang H, Guo S, Qiao X, Guo J, Li Z, Zhou Y, Bai S, Gao Z, Wang D, Wang P, et al. 2019. BZU2/ZmMUTE controls symmetrical division of guard mother cell and specifies neighbor cell fate in maize (S Hake, Ed.). PLOS Genetics 15: e1008377.

Weigel D, Glazebrook J. 2006. Setting up Arabidopsis crosses. CSH Protocols 2006: pdb.prot4623-pdb.prot4623.

Wu Z, Chen L, Yu Q, Zhou W, Gou X, Li J, Hou S. 2019. Multiple transcriptional factors control stomata development in rice. New Phytologist 223: 220–232.

Yang M, Sack FD, Altmann T. 1995. The too many mouths and four lips mutations affect stomatal production in Arabidopsis. The Plant Cell 7: 2227–2239.

Zhang D, Abrash EB, Nunes TDG, Prados IH, Gil MXA, Jesenofsky B, Lindner H, Bergmann DC, Raissig MT. 2022. Opposite polarity programs regulate asymmetric subsidiary cell divisions in grasses. bioRxiv: 2022.04.24.489281.

