## Supplementary material for "Extracurricular roles and divergent regulation of FAMA in *Brachypodium* and *Arabidopsis* stomatal development": Figs S1-10 and Tables S1-2

**Supplemental PDF File contains:**

Figure S1. Characterization of *BdFAMA* mutations and mutant phenotypes.

Figure S2. Additional phenotypes of *bdmute; BdFAMApro::YFP-BdFAMA* and *bdmute; BdMUTEpro::YFP-BdFAMA.*

Figure S3. Dose-dependent phenotypes of *bdmute; BdMUTEpro::YFP-BdFAMA* and *WT; BdMUTEpro::YFP-BdFAMA.*

Figure S4. Expression patterns of Brachypodium stomatal bHLH transgenes used in Co-IP experiments.

Figure S5. Bait enrichment in the Co-IP.

Figure S7. Summary of unique and shared of putative interaction partners from Co-IPs of stomatal bHLHs.

Figure S8. Y2H confirms bHLH interactions identified by Co-IPs.

Figure S9. Protein sequence alignments of MUTE and FAMA in Brachypodium and Arabidopsis.

Figure S10. Additional phenotypes revealed in rescue experiments with Arabidopsis and Brachypodium stomatal bHLHs.

Table S1. Primers sequences

Table S2. Summary of lines used

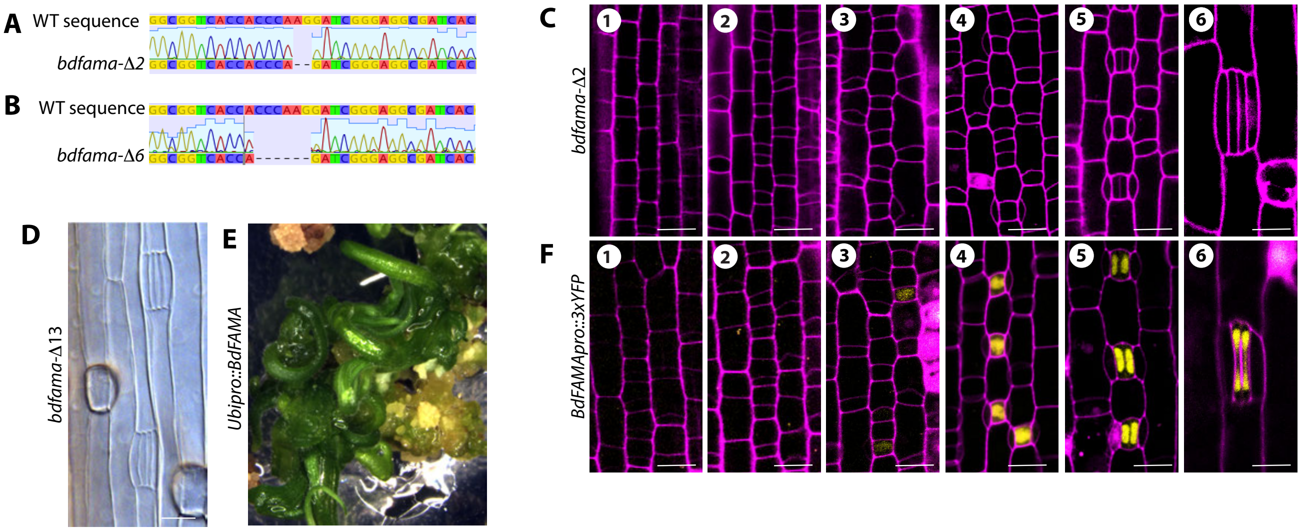

**Figure S1. Characterization of *BdFAMA* mutations and mutant phenotypes. (A-B)** Chromatograms of *bdfama-Δ2* (null mutant) and *bdfama-Δ6* (in-frame deletion with wild-type phenotype). **(C)** Confocal images of developmental stages of *bdfama-Δ2* during stomatal file establishment (stage 1), asymmetric division (stage 2), SMC establishment (stage 3), SC recruitment (stage 4), GC division (stage 5), and mature stomatal complexes (stage 6). **(D)** DIC images of cleared tissue from homozygous mutant T0 regenerant with 13bp deletion, *bdfama-Δ13.* **(E)** Image of regenerants from transformation with *Ubipro::BdFAMA* showing severe morphological defects in leaf tissues (5x magnification). **(F)** Confocal images of BdFAMA transcriptional reporter, *BdFAMApro::3xYFP,* during development. Cell outlines (magenta) visualized by Propidum Iodide (PI) staining. Scale bar=10 μm.

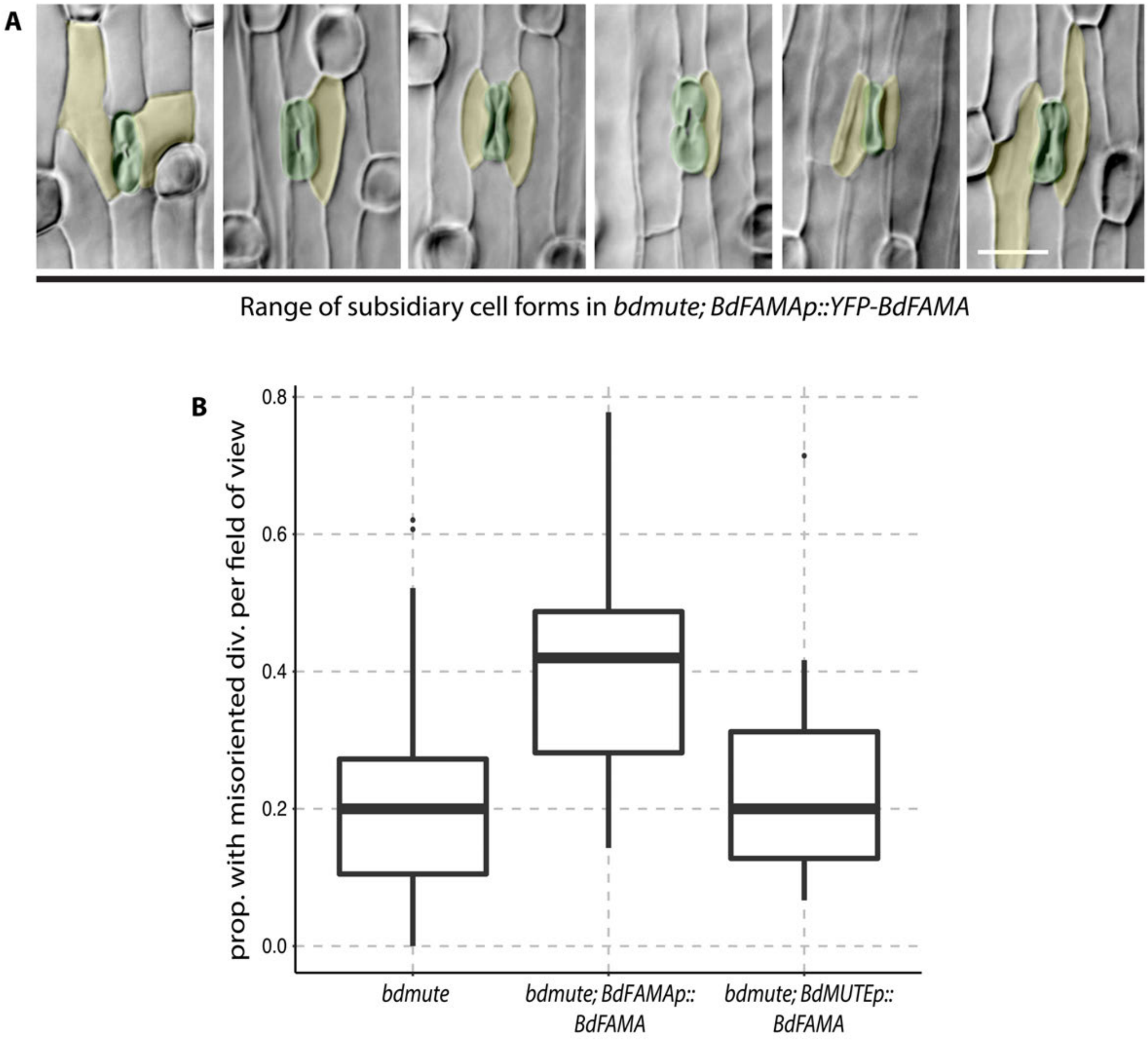

**Figure S2. Additional phenotypes of *bdmute; BdFAMApro::YFP-BdFAMA* and *bdmute; BdMUTEpro::YFP-BdFAMA.*** **(A)** Representative images of the phenotypic range of subsidiary cells recruited in *bdmute; BdFAMApro::YFP-BdFAMA*. Guard cells and presumptive subsidiary cells and psuedo-colored in green and yellow, respectively. **(B)** Proportion of stomatal complexes with misoriented GMC divisions per field of view in *bdmute, bdmute; BdFAMApro::YFP-BdFAMA* and *bdmute; BdMUTEpro::YFP-BdFAMa*. n=623, 2,354, and 1,349 stomata for *bdmute, bdmute; BdFAMApro::YFP-BdFAMA* and *bdmute; BdMUTEpro::YFP-BdFAMA,* respectively.

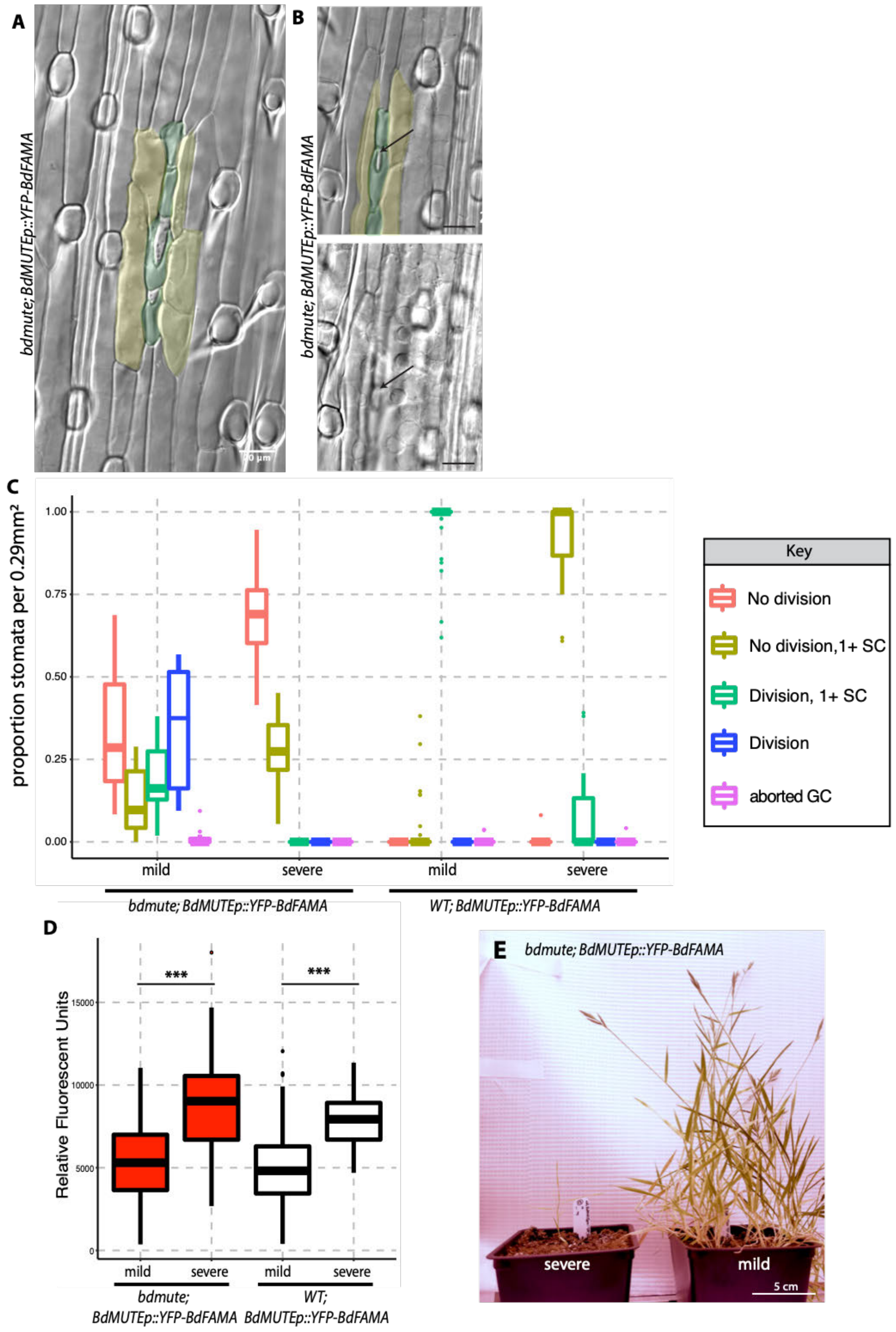

**Figure S3. Dose-dependent phenotypes of *bdmute; BdMUTEpro::YFP-BdFAMA* and *WT; BdMUTEpro::YFP-BdFAMA.*** **(A-B)** DIC images of stomatal cell clusters in *bdmute; BdMUTEpro::YFP-BdFAMA* severe lines that give rise to a pore. **(B)** Representative image of stomatal clusters of single guard cells that give rise to a pore with underlying mesophyll airspace. Cleared tissue from 2$^{rd}$ leaf, 6 to7 days post germination in T1 *bdmute; BdMUTEpro::YFP-BdFAMA* plants. **(C)** Quantification of stomatal phenotypes in *bdmute* and wild type (Bd21-3) lines expressing *BdMUTEpro::YFP-BdFAMA* from cleared abaxial tissue of the 2$^{rd}$ leaf, 6 to7 days post germination in T1 plants (dpg). Key (right) shows colors corresponding to phenotypic classes. For each sample, five different regions of the leaf were imaged and quantified; the same quantification of phenotypes used to generate Fig. 2C, *bdmute; BdMUTEpro::YFP-BdFAMA,* is used here, but stratified by line severity. n=1,349 stomata and 40 fields of view for *bdmute; BdMUTEpro::YFP-BdFAMA* and n=1,450 stomata and 47 fields of view for *WT; BdMUTEpro::YFP-BdFAMA*. **(C)** Quantification of relative fluorescence in severe and mild lines of *bdmute; BdMUTEpro::YFP-BdFAMA* (red boxes) and *WT; BdMUTEpro::YFP-BdFAMA* (white boxes) using confocal images from the same leaves used in (A). To standardize quantification, only cells in the developmental zone that fell between 4-6 µm in length and width were used in the analysis, n=130 cells for *bdmute; BdMUTEpro::YFP-BdFAMA* and n=172 cells for *WT; BdMUTEpro::YFP-BdFAMA*. ***P<0.001 (based on Wilcoxon-rank sum test followed by Dunn's multiple comparisons test). **(E)** Plant growth difference between mild and severe phenotypes of *bdmute; BdMUTEpro::YFP-BdFAMA.* Scale bar= 5cm.

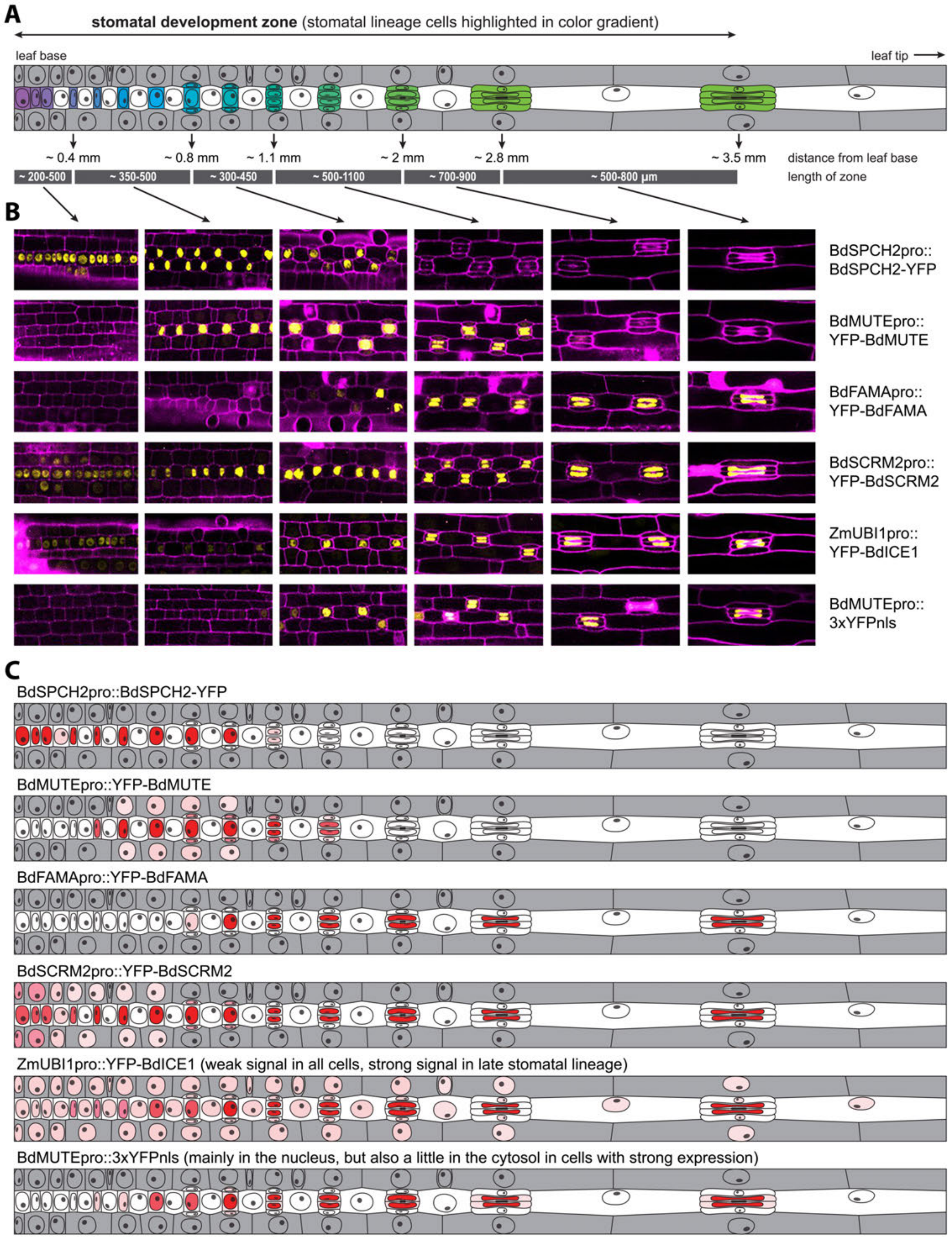

**Figure S4. Expression patterns of Brachypodium stomatal bHLH transgenes used in Co-IP experiments**
(A) Scheme showing the stages of stomatal development along the base of a young leaf. Stomatal lineage cells are highlighted with a purple-blue-green gradient. The approximate distance of each stage (first visible asymmetric cell

divisions, recruitment of subsidiary cells, symmetric division of the GMC, elongation of the complex, formation of the typical dumbbell shape of GCs, complex fully elongated) from the leaf base, as well as the length of each zone as determined by confocal microscopy of different plant lines is given below. (B) Confocal images showing expression of indicated reporter constructs in each zone in the lines used for the experiment (yellow). Leaves were stained with PI to visualize cell outlines (magenta). (C) For better comparison, expression of the reporter constructs in the epidermis is also shown in the schemes on the bottom with shades of red indicating the expression level within a line (red = high, pink = low). All signals were nuclear.

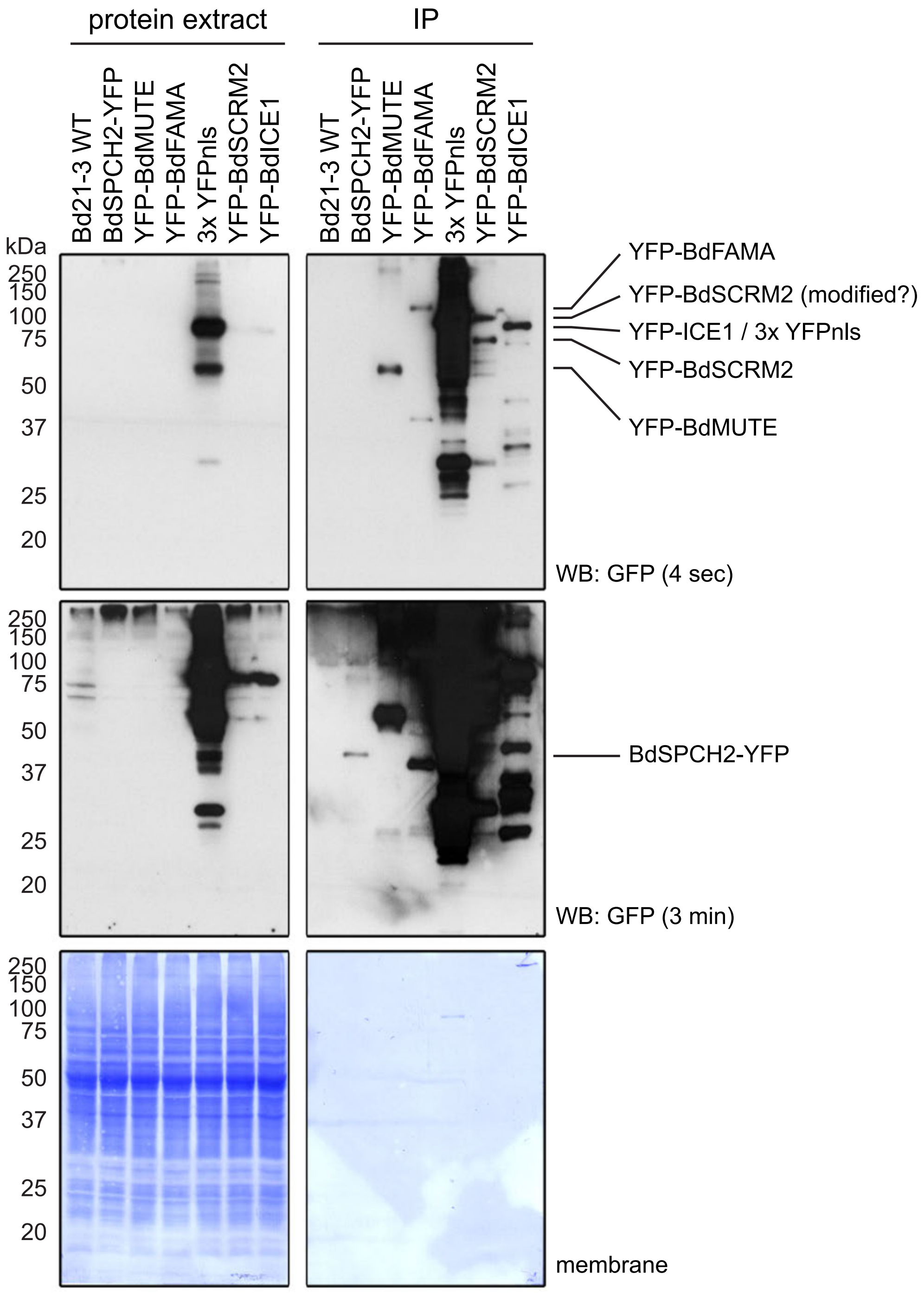

**Figure S5. Bait enrichment in the Co-IP**

Western blot (WB) analysis of proteins extracts before and proteins eluted from the beads after IP using an anti-GFP antibody. A short (top) and long (middle) exposure, as well as the Coomassie Brilliant Blue-stained membranes (bottom) are shown. The position of the YFP fusion proteins is indicated on the right.

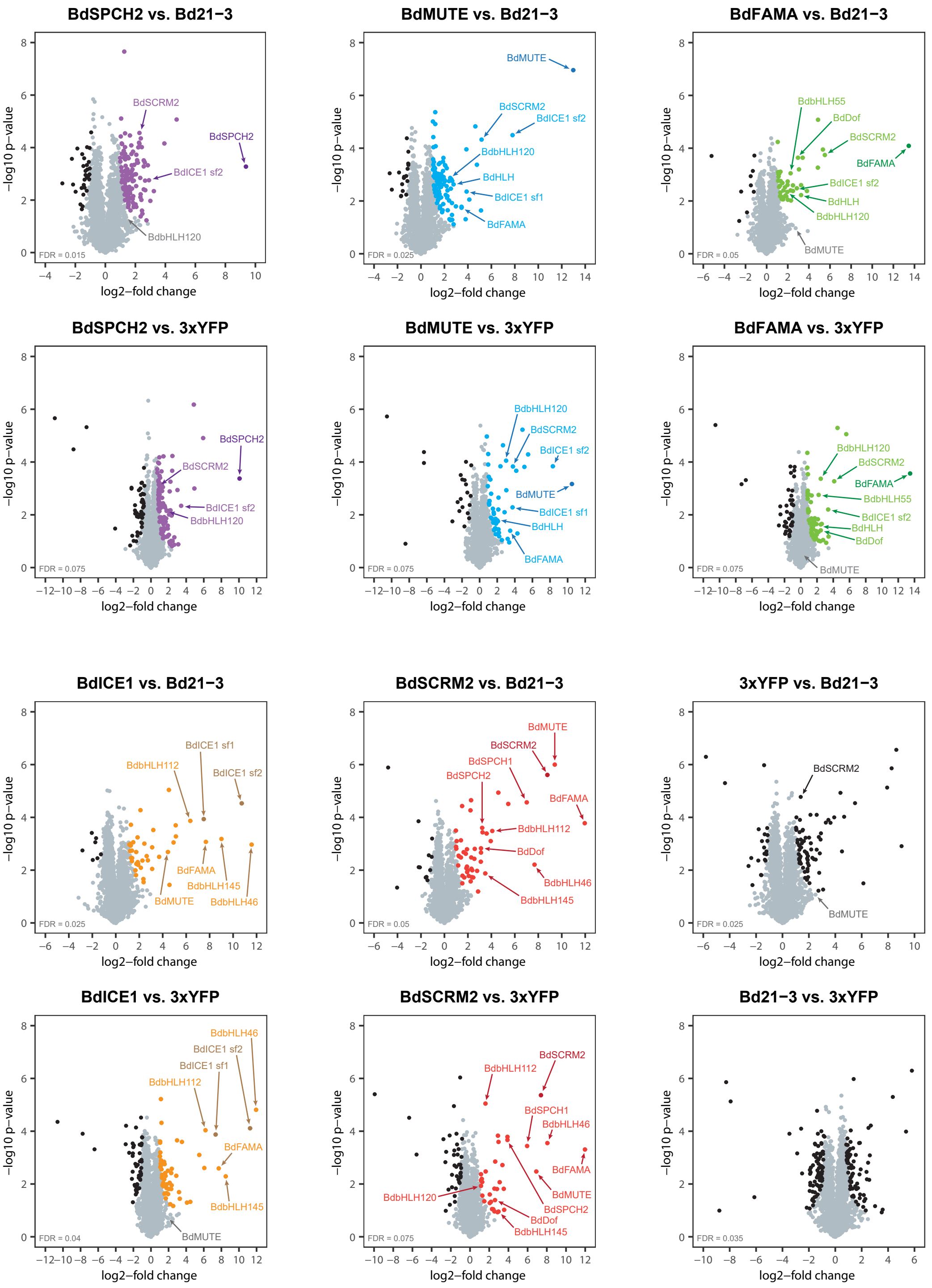

**Figure S6. Summary of proteins enriched in each stomatal bHLH co-IP experiment.** Scatter plots show the log2-fold change and -log10 p-value from unpaired 2-sided t-tests between the bHLH reporter lines and each of the two controls with a permutation-based FDR for multiple sample correction (S0 = 0.5). The FDR for each comparison was selected to minimize false negatives (proteins enriched in the control, highlighted in black). Proteins significantly enriched in the bHLH reporter lines are highlighted in color and the positions of candidates highlighted in this paper are indicated in the plots.

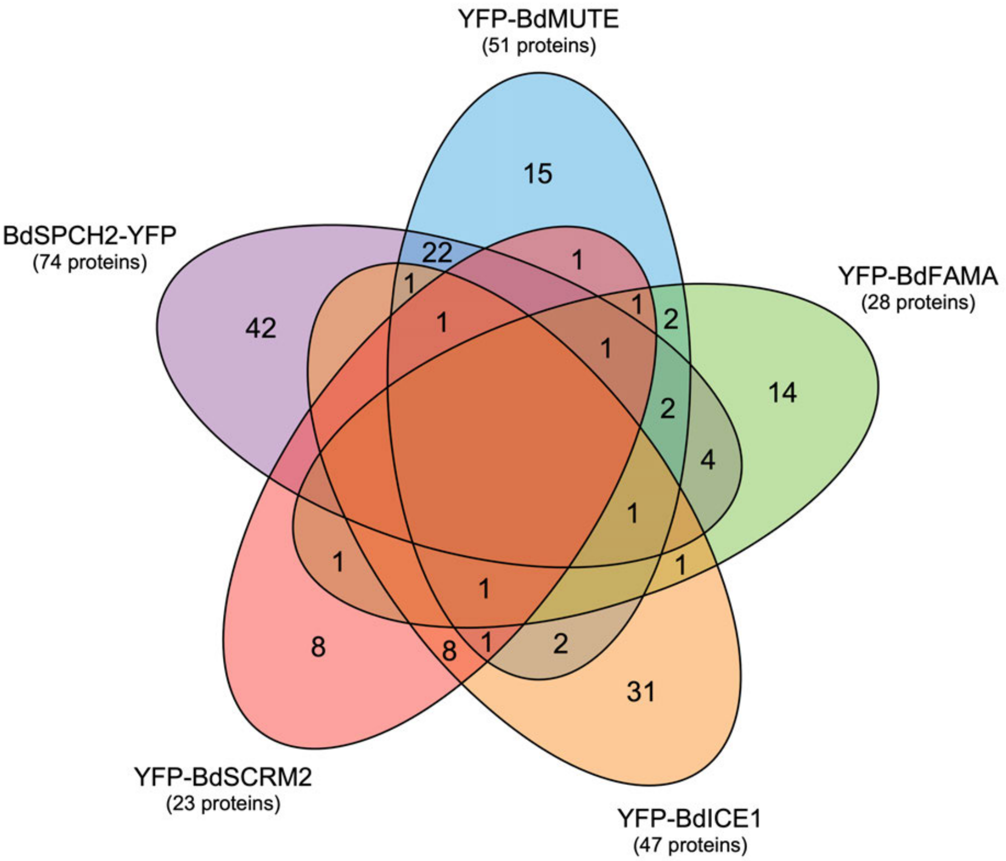

**Figure S7. Summary of unique and shared putative interaction partners from Co-IPs of stomatal bHLHs**
5-way Venn diagram showing the overlap of putative interaction partners of BdSPCH2 (purple), BdMUTE (blue), BdFAMA (green), BdICE1 (orange) and BdSCRM2 (red) identified by Co-IP and LC-MS/MS analysis.

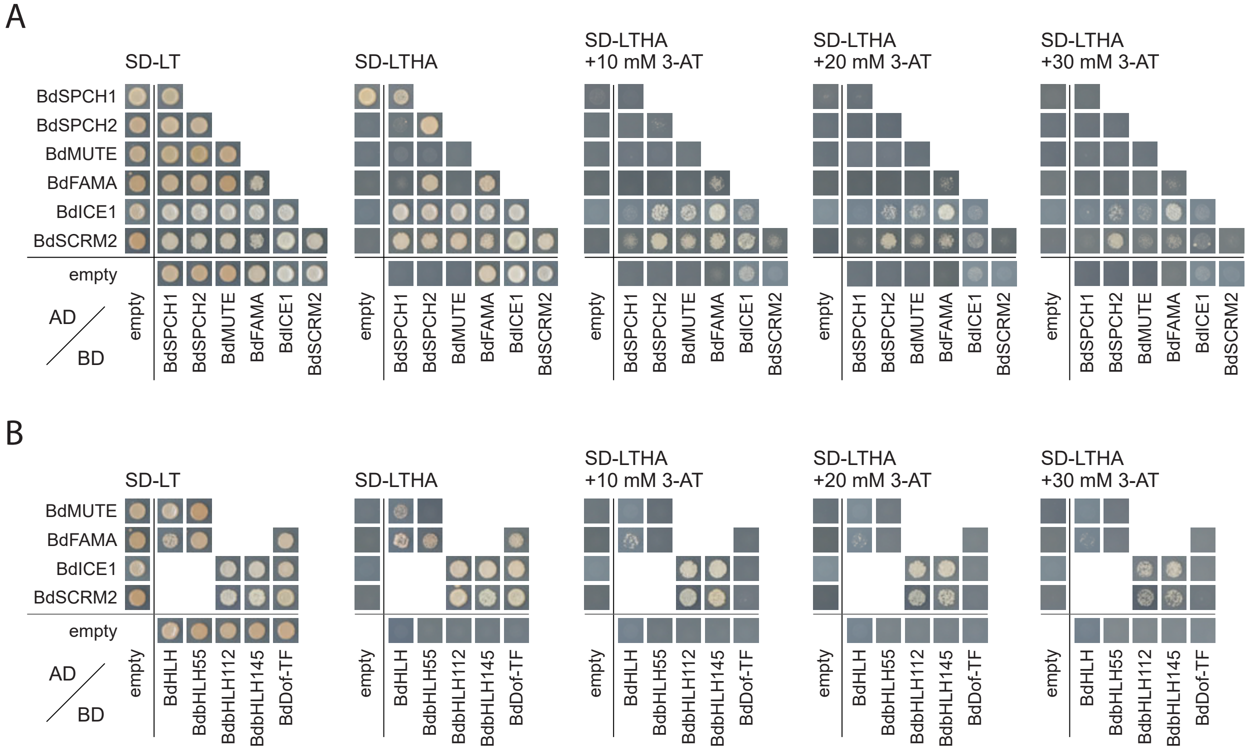

**Figure S8. Y2H confirms bHLH interactions identified by Co-IPs**
Y2H assay testing interactions between the six master regulators of stomatal development **(A)** and with selected additional TFs identified in the Co-IPs **(B)**. Yeast transfected with the indicated AD- and BD-fusion proteins was spotted on SD-LT and SD-LTHA to test for successful transfection and protein interaction, respectively. Different concentrations of 3-AT were added to increase stringency of the selection and to overcome auto-activation of BD fusions of BdFAMA, BdICE1 and BdSCRM2. Shown is one replicate of OD = 1 suspension. Controls with the empty BD plasmid are replicated in (A) and (B).

### Alignment of MUTE and FAMA in Arabidopsis and Brachypodium

```
AT3G24140-AtFAMA      -----MDKDYSAPNFLGESSGGNDDNSSGMIDYMFNRNLQQQQKQSMPQQQQHQLSPSGF
Bradi2g22810-BdFAMA   MEKQSEQGSNNQQQQQLDSFAPLDGAAPDQDQIIGGGAGAEMVDYMLGQQTPPPPPPPHG
AT3G06120-AtMUTE      ------------------------------------------------------------
Bradi1g18400-BdMUTE   ------------------------------------------------------------

AT3G24140-AtFAMA      GATPFDKMNFSDVMQFADFGSKLALNQTRNQDDQ—G--IDPVYFLKFPVLNDKIEDHNQ
Bradi2g22810-BdFAMA   HVSSFDKLSFSDVLHFADFGPRLALNQPLSTHHPADSDNDEDSYFFRFQPSLPAAEDSDD
AT3G06120-AtMUTE      ------------------------------------------------------------
Bradi1g18400-BdMUTE   ------------------------------------------------------------

AT3G24140-AtFAMA      TQHLMPSHQTSQEGGECGGNIGNVFLEEKEDQDDDNDNNSVQLRFIGGEEEDRENKNVTK
Bradi2g22810-BdFAMA   PTAQHAAVTTQGSGGDHGTVGGGVSESTTTLVQPQQQQQETVGGGKGGGGGGAGNK----
AT3G06120-AtMUTE      ------------------------------------------------------------
Bradi1g18400-BdMUTE   ------------------------------------------------------------

AT3G24140-AtFAMA      KEVKSKRKRARTSKTSEEVESQRMTHIAVERNRRKQMNEHLRVLRSLMPGSYVQRGDQAS
Bradi2g22810-BdFAMA   ---SGRRKRPRSTKTSEEVESQRMTHIAVERNRRRQMNDYLRVLRSLMPGSYVQRGDQAS
AT3G06120-AtMUTE      -----------------------MSHIAVERNRRRQMNEHLKSLRSLTPCFYIKRGDQAS
Bradi1g18400-BdMUTE   -----------------------MSHIAVERNRRRQMNEHLKTLRSLTPALYVKRGDQAS
                                             *:*********:***::*: **** *  *::******

AT3G24140-AtFAMA      IIGGAIEFVRELEQLLQCLESQKRRRILGETGRDMTTTTTSSSSPITTVAN-------QA
Bradi2g22810-BdFAMA   IIGGAIEFIRELEQLIQCLESQKRRRLYGDAPRPTAPDISTGAGAPPVVPP-------PA
AT3G06120-AtMUTE      IIGGVIEFIKELQQLVQVLESKKRRKTLN------------RPSFPYDHQTIEPSSLGAA
Bradi1g18400-BdMUTE   IIGGAVDFIRELHVLLEALQANKRRRLNNNLHPCSTPTTPSPRSLPTNNTNSSSPGSGGS
                      ****.::*::**. *:: *:::***:  .                .             :

AT3G24140-AtFAMA      QPLIITGNVTELE-------------------------------------------GGGG
Bradi2g22810-BdFAMA   TSSMLQHEQQAAPPQGPPHHDAPAPFYVVPAPSPGTSLPLIPVISDDGVAKGIDDLDGGL
AT3G06120-AtMUTE      TTRVPFSRIE-------------------------------------------NVMTTST
Bradi1g18400-BdMUTE   SSAASNTGSG-------------------------------------------GGVNKEK
                      .

AT3G24140-AtFAMA      LREETAENKSCLADVEVKLLGFDAMIKILS-RRRPGQLIKTIAALEDLHLSILHTNITTM
Bradi2g22810-BdFAMA   GREEVAENKSCLADIEVRVLGADAVVKVLS-RRRPEQLIKTIAVLEEMHLSILHTNITTI
AT3G06120-AtMUTE      FKEVGACCNSPHANVEAKISGSNVVLRVVS-RRIVGQLVKIISVLEKLSFQVLHLNISSM
Bradi1g18400-BdMUTE   ARELAACCSSAAAEVEARISGANLLLRTLSGRAPPGQAAKMVGLLQALHLEVLHLNISTL
                      :*  *  .*  *::*.:: * : ::: :* *      *  * :. *: : :.:** **::

AT3G24140-AtFAMA      EQTVLYSFNVKITSETRFTAEDIASSIQQIFSFIHANTNISGSSNLGNIVFT--------
Bradi2g22810-BdFAMA   DQTVLYSFNVKIAGEPRFTAEDIAGAVHQILSFIDINYTL--------------------
AT3G06120-AtMUTE      EETVLYFFVVKIGLECHLSLEELTLEVQKSFVSDEVIVSTN-------------------
Bradi1g18400-BdMUTE   EDTVLHSFVLQIGLECQLSVEDLAFEVHQTFCCDYQQEDHHGQQQLLLELPIAGTVHGDI
                      ::***: * ::*  * ::: *:::  ::: :

AT3G24140-AtFAMA      ---
Bradi2g22810-BdFAMA   ---
AT3G06120-AtMUTE      ---
Bradi1g18400-BdMUTE   MIN
```

**Key**

bHLH domain | LxCxE binding site
HER DNA binding site | MUTE SMF domain
FAMA SMF domain

**Figure S9. Protein sequence alignments of MUTE and FAMA in Brachypodium and Arabidopsis**
Protein alignment of *AtMUTE, AtFAMA, BdMUTE,* and *BdFAMA.* The bHLH domain is indicated by blue text, and contains the HER DNA binding domain (pink highlight). The LxCxE RBR binding site of AtFAMA, and IxCxE alternative site of BdFAMA are shown in yellow highlight. The ACT-like SPCH-MUTE-FAMA (SMF) structural domain is highlighted in blue. Alignments were done using Clustal 2.1 multiple sequence alignment tool.

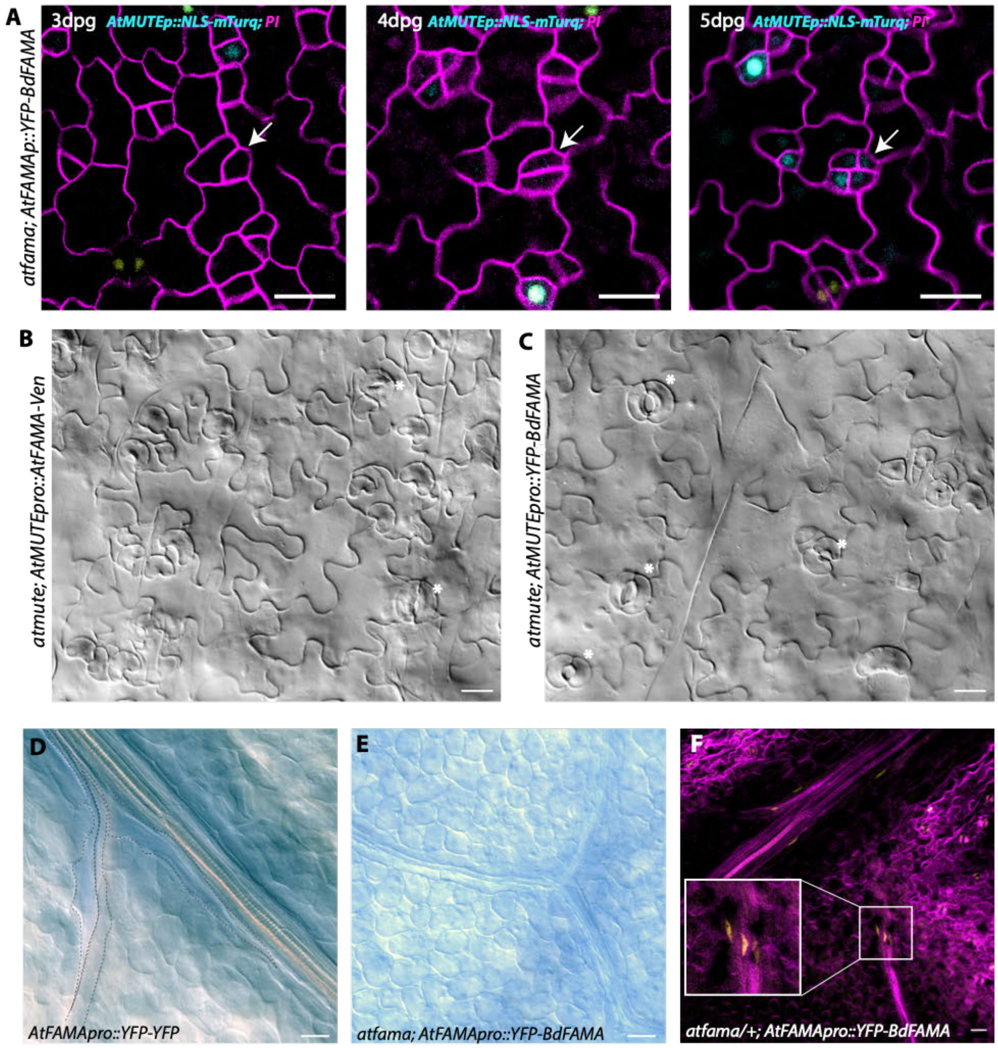

**Figure S10. Additional phenotypes revealed in rescue experiments with Arabidopsis and Brachypodium stomatal bHLHs. (A)** Confocal images taken in time course at 3, 4, and 5 dpg in *atfama; AtFAMApro::YFP-BdFAMA; AtMUTEpro::NLS-mTurq*. White arrow points to a cell at 3dpg that divides and expresses *AtMUTE* at low levels at 4dpg, and divides again at 5dpg, with all daughter cells expressing *AtMUTE*. **(B-C)** DIC images of cleared abaxial tissue from *atmute; AtMUTEpro::AtFAMA-Ven* and *atmute; AtMUTEpro::YFP-BdFAMA* in 10dpg cotyledons. White asterisk indicates stoma. **(D-E)** DIC images of tissue stained with Coomassie Brilliant Blue (CBB) to visualize the presence or absence of myrosin idioblast (MI) cells in *AtFAMApro::YFP-YFP* and *atfama; AtFAMApro::YFP-BdFAMA*. **(D)** Developing MIs are stained blue in *AtFAMApro::YFP-YFP*. **(E)** *AtFAMApro::YFP-BdFAMA* fails to rescue the MI development defect in *atfama* and no stained MIs are observed. **(F)** Confocal image of abaxial tissue from true leaf *atfama/+; AtFAMApro::YFP-BdFAMA* stained with PI. Signal is observed in the heterozygous *atfama* background. All myrosin cell images are taken from first rosette true leaves 10-15 dpg. Scale bar=20 µm.

**Table S1. Primer sequences**

| Primer name | Sequence | Purpose | Notes |
|---|---|---|---|
| primMXA 22-FWD | GCTATGATCTCTCGCAGTCG | Genotyping BdFAMA CRISPR g4 | |
| primMXA 24-REV | AACGGAGAGAGAGTACTACGGG | Genotyping BdFAMA CRISPR g4 | |
| priJM_BdMUTEpro-FWD | CAGGCTAGCAGCACTATT | Genotyping BdMUTE promoter for cloning | from Raissig et al 2017 |
| priJM_BdMUTEpro-REV | GATCGTGTCGTTCTTCct | Genotyping BdMUTE promoter for cloning | from Raissig et al 2017 |
| priDZ44, Hyg_MRaissig_Fprim | GCGAGTACTTCTACACAGCC | Genotyping Hygromycin resistance in Brachypodium lines | from Raissig et al 2016 |
| priDZ45, Hyg_MRaissig_Rprim | GCGAAGAATCTCGTGCTTTC | Genotyping Hygromycin resistance in Brachypodium lines | from Raissig et al 2016 |
| priDZ46, Cas9_MRaissig_Fprim | TCGACGAACAGCTGCTTTT | Genotyping Cas9 in Brachypodium lines | from Raissig et al 2016 |
| priDZ47, Cas9_MRaissig_Rprim | GGACAAGGGCAGGGATTTC | Genotyping Cas9 in Brachypodium lines | from Raissig et al 2016 |
| BdSCRM2 qPCR-FWD | GCCGAGCAATGGAAGGATGGTC | qPCR primer for BdSCRM2-forward | |
| BdSCRM2 qPCR-REV | AGACTAAGGGCCACAGTTCCGG | qPCR primer for BdSCRM2-reverse | |
| BdMUTE qPCR-FWD | TCGAAGCACTCCAGGCAAACAAG | qPCR primer for BdMUTE-forward | |
| BdMUTE qPCR-REV | GGTGCTGCAGGGATGAAGATTG | qPCR primer for BdMUTE-reverse | |
| BdFAMA qPCR-FWD | TACTCCTTCAACGTCAAGATCGC | qPCR primer for BdFAMA-forward | |
| BdFAMA qPCR-REV | GTCGATGAAGCTAAGGATCTGGTG | qPCR primer for BdFAMA-reverse | |
| BdICE1 qPCR-FWD | TGGATGTCTTCAAGGCTGAGGTAG | qPCR primer for BdICE1-forward | from Raissig et al 2016 |
| BdICE1 qPCR-REV | GACTTGGAGCAGAACTGCCTTG | qPCR primer for BdICE1-reverse | from Raissig et al 2016 |
| priMR312 | GTCACCCGCAATGACTGTAAGTTC | qPCR primer for BdUBC18 - Bradi4g00660 | from Raissig et al 2016 |
| priMR313 | TTGTCTTGCGGACGTTGCTTTG | qPCR primer for BdUBC18 - Bradi4g00660 | from Raissig et al 2016 |
| priMXA3 | ggcaGGCGGTCACCACCCAAGGAT | Guide 4 CRISPR Bradi2g22810 BdFAMA FP | |
| priMXA4 | aaacATCCTTGGGTGGTGACCGCC | Guide 4 CRISPR Bradi2g22810 BdFAMA RP | |
| priMXA5 | ATGGAAAAACAGGTCTGCTT | Cloning BdFAMA ORF | |
| priMXA6 | CCTCCATGAGAAAGTGGAAG | Cloning BdFAMA ORF with 3' UTR | |
| MUTE SEQ F | ATGTCTCACATCGCTGTTGAAAGGAATCG | Genotyping *atmute* | |
| MUTE SEQ R | ATCGAAGCTTGATCTCCCTAATACCGATC | Genotyping *atmute* | |
| primMXA7 | GGCGCGCCACCAGCCTAGGAGAGTTGAG | Cloning BdFAMA promoter | |
| primMXA8 | GGCGCGCCTGATCAGAAGGAACACGTATGG | Cloning BdFAMA promoter | |

| | | |
|---|---|---|
| gAtFAMA-fw | CACCATGGATAAAGATTACTCGGTACGTACG | cloning AtFAMA gDNA FWD |
| gAtFAMA-rev | AGTAAACACAATATTTCCCAGGTTAGAGC | cloning AtFAMA gDNA REV |
| fama-1 LP | TCATTCATTTGCTTCCTACGG | Genotyping atfama-1 TDNA Salk line (Salk_100073) |
| fama-1 RP | CAATACAAAAAGCTCCCCTCAC | Genotyping atfama-1 TDNA Salk line (Salk_100073) |
| LBb1 | GCGTGGACCGCTTGCTGCAACT | Genotyping atfama-1 TDNA Salk line (Salk_100073) |
| BdSPCH1_fw | CACCATGGGAGACATCGCGCTGT | Cloning of Bradi1g38650 |
| BdSPCH1_rev | TCACGAGAACGTTTGCTGAATCTCT | Cloning of Bradi1g38650 |
| BdSPCH2_fw | CACCATGGCCATGGGGGATGAC | Cloning of Bradi3g09670 |
| BdSPCH2_rev | TCACAAAAAGGTCTGCCGGA | Cloning of Bradi3g09670 |
| BdMUTE_fw | CACCATGTCGCACATCGC | Cloning of Bradi1g18400 |
| BdMUTE_rev | TTAATTGATCATGATGTCGC | Cloning of Bradi1g18400 |
| BdFAMA_fw | CACCATGGAAAAACAGTCGGAGCAG | Cloning of Bradi2g22810 |
| BdFAMA_rev | TCATAACGTGTAGTTGATGTCGATG | Cloning of Bradi2g22810 |
| BdICE1-sf1_fw | CACCATGCTGTCGGGGTTCAAC | Cloning of Bradi4g17460.1 |
| BdICE1-sf1_rev | CTAGATCATCGGATGGAACCC | Cloning of Bradi4g17460.1 |
| BdSCRM2_fw | CACCATGGAGAATTCGGTGGGGG | Cloning of Bradi2g59497 |
| BdSCRM2_rev | CTACATTGGGTTCTGAAGACCGG | Cloning of Bradi2g59497 |
| BdHLH_fw | CACCATGATGTCGAGGGAGCG | Cloning of Bradi1g63040 |
| BdHLH_rev | CTATATTTGCTCGTCGTCGTG | Cloning of Bradi1g63040 |
| BdbHLH55_fw | CACCATGGTGATGAAGATGGAGGTAGAG | Cloning of Bradi2g08080 |
| BdbHLH55_rev | CTACACCATGTTCATGGCACG | Cloning of Bradi2g08080 |
| BdbHLH112_fw | CACCATGGCACTAGTGGACGCG | Cloning of Bradi3g52150 |
| BdbHLH112_rev | TTACTGCGAGGCCACGAG | Cloning of Bradi3g52150 |
| BdbHLH145_fw | CACCATGACGCTGGATGCCGT | Cloning of Bradi5g19950 |
| BdbHLH145_rev | TTAACTTAAAAGTTCCGGCTGCC | Cloning of Bradi5g19950 |
| BdDof_fw | CACCATGATTCCCATCGATCTCCAA | Cloning of Bradi1g15420 |
| BdDof_rev | CCCTAGGGCAGCATGG | Cloning of Bradi1g15420 |

### Table S2. Summary of lines used

| Plant line | Plant species | Construct used | Background genotype | Generation imaged in paper | # T0 regenerants | # Regenerants that lived to produce seeds |
|---|---|---|---|---|---|---|
| *bdfama* | *Brachypodium distachyon* | pEX of BdFAMA Guide 4 - sgRNA under Ubip::Cas9 | Bd21-3 | T3 | 4 | 3; 2 with out of frame mutation, 1 in frame |
| *BdFAMApro::YFP-BdFAMA* | *Brachypodium distachyon* | pEX_BdFAMApro::BdFAMA-YFP | Bd21-3 | T3 | 15 | 13 |
| *BdFAMApro::3xYFPnls* | *Brachypodium distachyon* | pEX_BdFAMApro::3xYFPnlsI | Bd21-3 | T1 | 19 | 13 |
| *UBIpro::BdFAMA* | *Brachypodium distachyon* | pEX_Ubipro::BdFAMA ORF | Bd21-3 | T0 | 8 | 2 (reporter silenced) had WT phenotype and no expression |
| *UBIpro::BdFAMA-YFP* | *Brachypodium distachyon* | pEX_Ubipro::BdFAMA-YFP | Bd21-3 | T0 | 4 | 1 |
| *bdmute/sid* | *Brachypodium distachyon* | EMS, bdmute-1 | *bdmute-1* | --- | --- | --- |
| *bdmute; BdFAMApro::YFP-BdFAMA* | *Brachypodium distachyon* | pEX_BdFAMApro::BdFAMA-YFP | *bdmute-1* | T3 | 60 (only imaged ~30) | 60 |
| *bdmute; BdMUTEpro::YFP-BdFAMA* | *Brachypodium distachyon* | pEX_BdMUTEpro::YFP-BdFAMA | *bdmute-1* | T0 and T1 | 19 | 5 |
| WT; *BdMUTEpro::YFP-BdFAMA* | *Brachypodium distachyon* | pEX_BdMUTEpro::YFP-BdFAMA | Bd21-3 | T0 and T1 | 17 | 8 |
| *bdmute; bdfama* | *Brachypodium distachyon* | pEX of BdFAMA Guide 4 - sgRNA under Ubip::Cas9 | *bdmute-1* | T0 | 28 | 0 |
| atmute/+ (G>A allele) | *Arabidopsis thaliana* | EMS, seed stock 686 | Col-0 | --- | | |
| atfama/+ (Salk T-DNA line) | *Arabidopsis thaliana* | Salk T-DNA line (Salk_100073) | Col-0 | --- | | |
| *atfama; AtFAMApro::YFP-BdFAMA* | *Arabidopsis thaliana* | pEX_AtFAMApro::YFP-BdFAMA | Col-0 | T3 | | |
| *atmute; AtMUTEpro::YFP-BdFAMA* | *Arabidopsis thaliana* | pEx_AtMUTEpro::YFP-BdFAMA | *atmute* | T3 | | |
| *atmute; AtMUTEpro::AtFAMA-Ven* | *Arabidopsis thaliana* | crossed with *atmute/+* | *atmute/+* | F3 | | |
| *atfama; AtFAMApro::AtFAMA-YFP* | *Arabidopsis thaliana* | R4GWB640-AtFAMApro::gAtFAMA-YFP | *atfama/+* | | | |
| *atfama; AtFAMApro::YFP-BdFAMA; AtFAMApro::NLS-mTurq* | *Arabidopsis thaliana* | pEX_AtFAMApro::NLS-mTurq | *atfama; AtFAMApro::YFP-BdFAMA* | T2 | | |
| *atfama; AtFAMApro::YFP-BdFAMA; AtMUTEpro::NLS-mTurq* | *Arabidopsis thaliana* | pEX_AtMUTEpro::NLS-mTurq | *atfama; AtFAMApro::YFP-BdFAMA* | T2 | | |
